# A Measure of Whole-Brain White Matter Topography

**DOI:** 10.1101/783274

**Authors:** Junyan Wang, Yonggang Shi

**Affiliations:** Laboratory of Neuro Imaging (LONI), USC Stevens Neuroimaging and Informatics Institute, Keck School of Medicine of USC, Los Angeles, CA 90033

## Abstract

The unprecedentedly high-quality large-scale brain imaging datasets, from such as the Human Connectome Project (HCP) and UK-Biobank, provide a unique opportunity for measuring the white matter topography of the human brain. By leveraging the multi-shell diffusion MRI data from the original young adult HCP, we systematically develop a reliable measure of the whole-brain white matter topography, and we coin it *topographic vector*. As the main result, we find that the three most dominant dimensions of the topographic vectors strongly and linearly correlate with the coordinates of the corresponding streamlines of the whole-brain tractograms. Our results support the earlier prescient hypothesis that brain development follows a “base-plan” established by three (main) chemotactic gradients of early embryogenesis, and they implicate that the whole brain white matter tracts can be represented by vectors of a natural coordinate system.

---

The topographic organisation, or topography, of the brain white matter was first measured using Klingler’s brain dissection technique in early post-mortem studies 30 years ago^1^. However, our understanding of it is still limited. We view the topography of brain white matter pathways at two scales: local and global. The local topographic organisation, which is also known as the topographic regularity, has been defined as *the preservation of the spatial relationship between neurons in point-to-point or region-to-region axonal connections*^*2,3*^ and it has been supported by tracer injection results^4-7^ and diffusion magnetic resonance imaging (dMRI) studies^8-11^ for various white matter pathways. According to its conceptual definition, a measure of the local topographic organisation of white matter streamlines has also been proposed very recently^12,13^. However, the global topography of the whole-brain white matter pathways is still barely understood, despite the persevering progress in this direction. For example, the anatomical organisation of the white matter pathways in macaque brains, as revealed by tracer injections, has been meticulously documented in a comprehensive monograph^5^. The topography of the cortico-striatal anatomical connectivity in macaque brains, as a core part of the brain reward network, and its association to incentive-based learning has been revealed^4,6,7^. More recently, based on dMRI tractography, geometrical characteristics of the white matter pathways in nonhuman primates and human brains have also been summarised in^14^. Additionally, based on their observations, the authors also made prescient hypotheses on the principle of human brain white matter development which reconcile with previous biological studies^15-17^. However, since there exists no effective method for us to measure the global topography of the whole-brain white matter tracts *in vivo*, the existing studies on it are mainly qualitative, which we believe to be a methodological obstacle to the precise understanding of its underpinning principle.

Thanks to the rapid development in brain imaging and the associated information technologies, we now have access to the large-scale unprecedentedly high-quality brain imaging datasets of various imaging modalities from, such as the Human Connectome Project^18,19^ and the UK-biobank^20^. Besides, the dMRI tractography is the only method for *in vivo* and non-invasive measurements of the white matter pathways of the human brains. These ground-breaking advancements grant us a unique opportunity for the development of novel computational methods for *in vivo* measuring the topography of the human brain white matter and beyond.

As the main contribution of this work, we present a quantitative study of the global topography of the whole-brain white matter tracts based on dMRI tractography. Our study is based on a novel multidimensional measure of the whole-brain white matter topography derived from a machine-learning technique called multidimensional scaling (MDS)^21,22^. The MDS maps the tractography streamlines to vectors in a finite-dimensional Euclidean vector space while preserving the pairwise distances between the streamlines in the mapping. The resultant measure naturally coincides with the biological notion of the global topography of white matter tracts in that, in mathematical terms, the metric geometry of the tractography streamlines is the same as that of the embedding vector set, and each dimension of the embedding vectors uniquely contributes to the metric geometry. Accordingly, we call the MDS embedding vectors the *topographic vectors*. In addition, to derive a reliable measure of the whole brain white matter topography, we systematically develop new scalable computational methods to extend the individual measure to a group-wise measure. The details of the methods can be found in the Methods section and the Supplementary Information.

In this work, we use the minimally preprocessed dMRI data from the original Human Connectome Project (HCP). We refer the readers to ^23^ for the details on the data acquisition and preprocessing. We refer to a set of tractography streamlines as a *tractogram* henceforth in this paper.

## Individual whole-brain streamline topography analysis

The topographic vectors for each individual tractogram are computed using MDS. We include the details of the computation in the Methods section. In **Figure 1**, we visualise the top-3 topographic vectors for the whole brain tractogram of an individual subject as a point cloud. A striking observation is that the enclosing shape of the topographic vectors resembles the shape of the brain. We can also observe certain internal structures of the point cloud.

**Figure 1:**
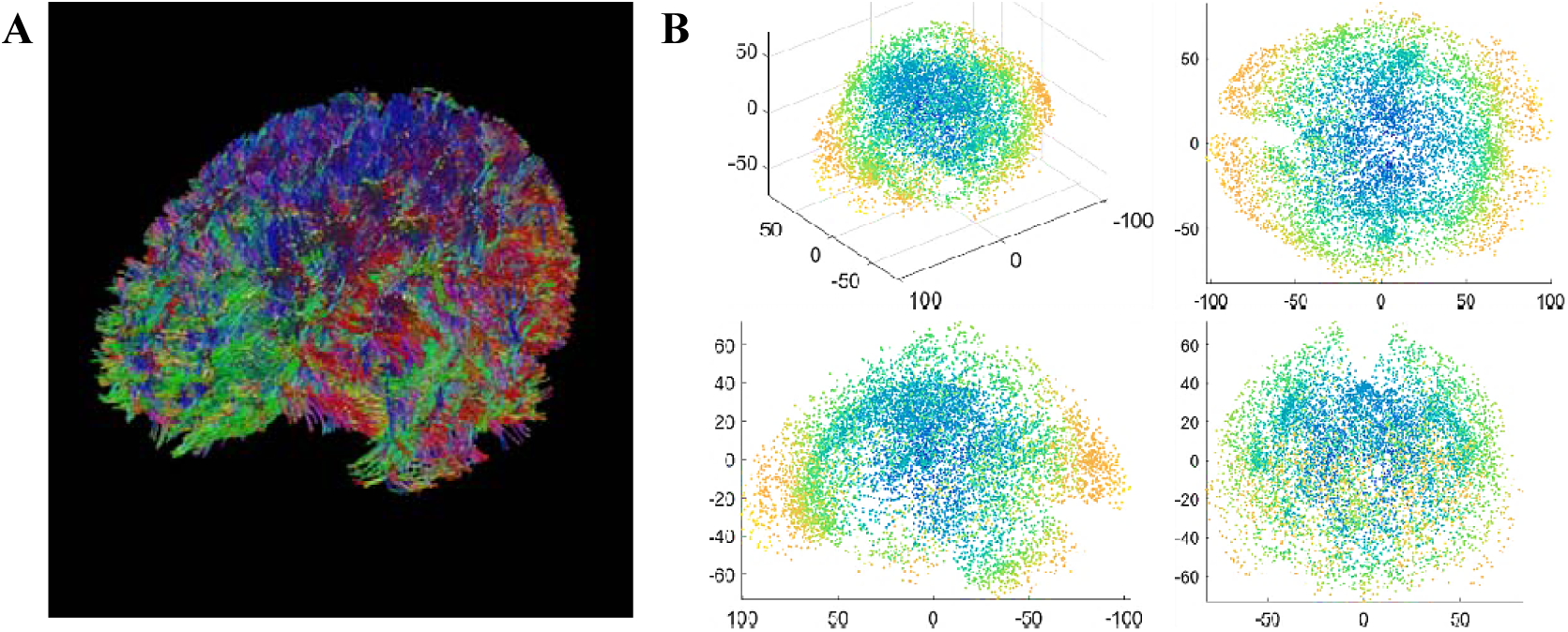
Topographic vector of an individual tractogram. A) is an example of a whole-brain tractogram visualised using TrackVis^a^. B) shows different views of the point cloud of the topographic vectors of the tractogram shown in (A).

Before analysing the topographic vectors, we examine the performance of the individual MDS embeddings across different subjects under three different settings: a) the tractograms of all subjects are in the native space of the T1-weighted MRI, b) the tractograms are warped to the MNI152 standard space based on volumetric T1-weighted image registration c) the tractograms are warped towards a reference subject using Track Density Image (TDI) ^24^ based volumetric registration. A total of 99 subjects were used in this experiment. According to the theory of MDS, the distance of the original space is Euclidean *iff* the eigenvalues of the Gram matrix are non-negative^21^. Hence, we are only concerned about the Euclidean part of the distance matrix. First of all, we evaluate the stability of the dimensions of embedding. The histograms of the maximum dimensions for the three sets of tractograms are shown in **Figure 2** (A), from which we can conclude that the tractograms are representable by roughly the same number of topographic vectors, and this suggests that the complexity of the tractograms is stable across different subjects.

**Figure 2:**
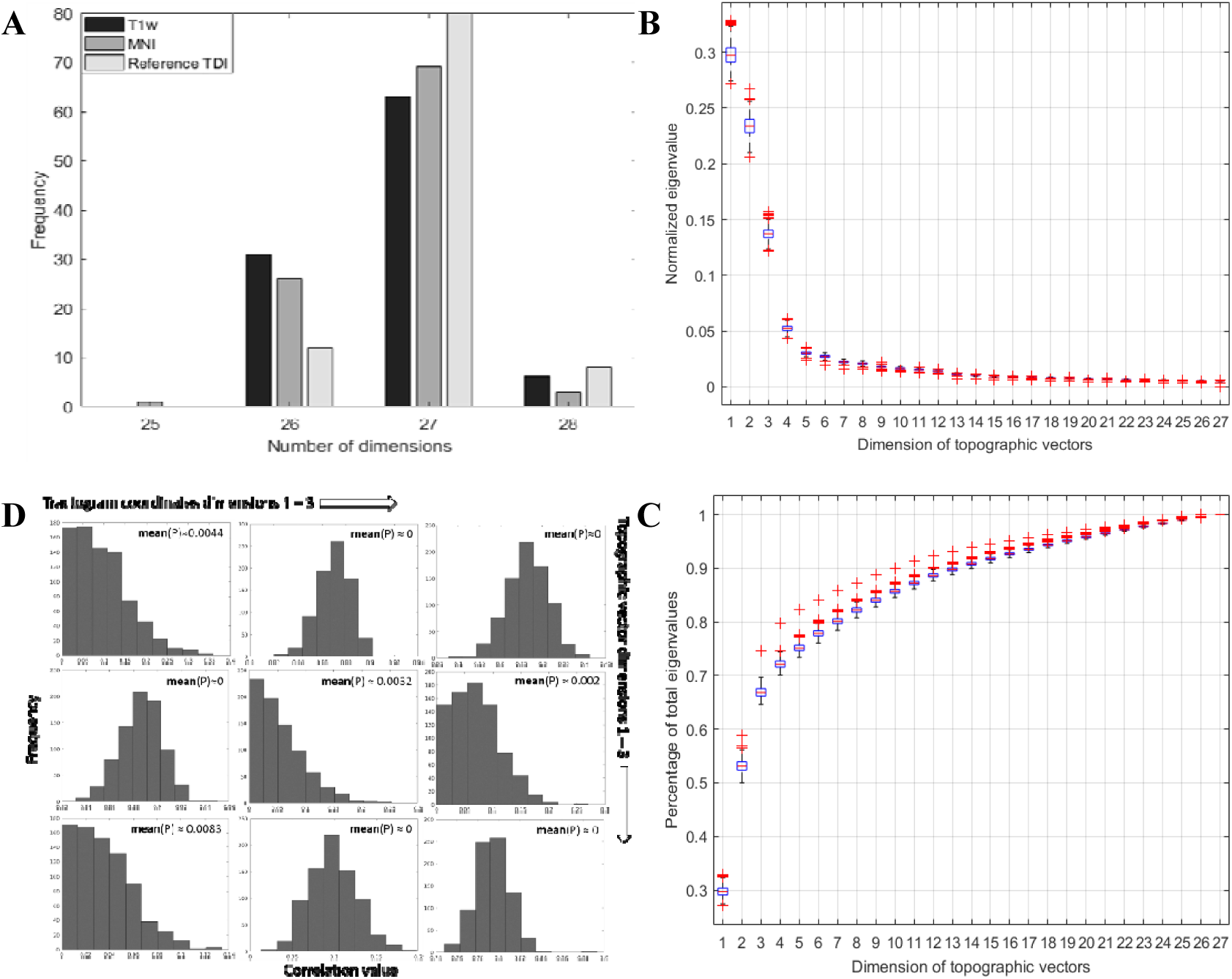
A) Histogram of maximum embedding dimensions for 10K streamline tractograms of HCP subjects under three settings. B) Boxplot of the eigenvalues of each dimension of the topographic vectors. C) Boxplots of the proportion of the first k positive eigenvalues over the total positive eigenvalues. D) Histograms of the linear correlations between the individual top-3 topographic vectors and the brain anatomical coordinates of the streamlines. The average p-values are also shown for each pairwise relation.

Afterwards, we evaluate the quality of the embedding in terms of distance preservability by calculating the correlation *ρ* between the pairwise distances of the topographic vectors and the input pairwise distance that we used to compute the embedding. The means and standard deviations of the correlations are summarized in **Table 1**. The quality of embedding is equally good in the three different settings, meaning that spatial normalization by image registration does not affect the quality of embedding. In addition, we can conclude that the distance metric of the original tractograms is approximately Euclidean.

**Table 1:**
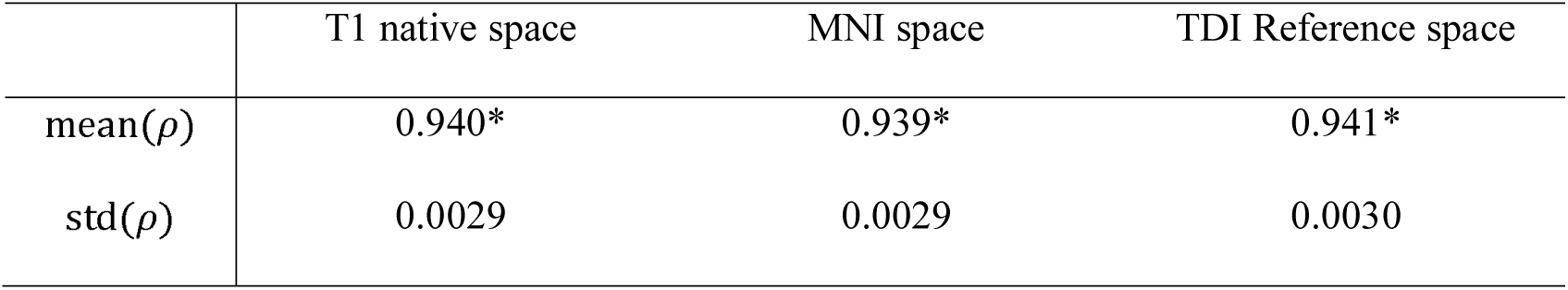
Quality of MDS embedding of whole-brain streamline bundles in three settings (* indicates statistical significance at α=0.05)

Since the contribution of each embedding dimension to the overall distance geometry is characterized by the corresponding eigenvalue, we proceed to study the eigenvalues of the MDS for each individual tractogram. The theoretical discussions on the eigenvalues of the embedding can be found in the Supplemental Information. In **Figure 2** (B) and (C), we show the boxplot of the distributions of the eigenvalues of all embedding dimension for all subjects in our study ^b^. The result shows that the eigenvalues are stable across different subjects, and the top-3 eigenvalues are significantly larger than the remaining dimensions.

In order to understand to what extend the topographic vectors represent the topography of the streamline curves, we compute the correlation between the topographic vectors of the tractograms in the TDI reference space and the corresponding streamline coordinates. 861 subjects were selected for this experiment. Details on the subject selection can be found in the Methods section. Since each topographic vector corresponds to one whole streamline curve, we may view the topographic vector as extra dimensions to the streamline coordinates. Accordingly, we can calculate the Pearson linear correlation between each dimension of the streamlines and each dimension of the topographic vectors. The histograms of the absolute values of the correlations are shown in **Figure 2** (D), from which we observe that each dimension of the top-3 topographic vectors strongly linearly correlates with one dimension of the streamline coordinates. Specifically, the 1^st^ dimension of the streamline coordinates strongly and linearly correlates with the 2^nd^ dimension of the topographic vectors, the 2^nd^ dimension of the streamline coordinates strongly and linearly correlates with the 1^st^ dimension of the topographic vectors, the 3^rd^ dimension of the streamline coordinates strongly and linearly correlates with the 3^rd^ dimension of the topographic vectors. As the coordinates of the streamlines are defined based on the anatomical axes of the brain, we would restate the conclusion as *the values of each of the top-3 dimensions of the topographic vectors strongly and linearly correlate with the streamline points in one of the 3 brain anatomical directions: anterior-posterior (1*^*st*^ *dimension), left-right (2*^*nd*^ *dimension) or dorsal-ventral (3*^*rd*^ *dimension).*

Small but significant correlations are found between the 2^nd^ dimension of streamline coordinates and the 3^rd^ dimension of the topographic vectors as well as the 1^st^ dimension of the topographic vectors and the 3^rd^ dimension streamline coordinates. In this experiment, no alignment across different subjects has been performed before the calculation of correlations. This experiment setting further strengthens our conclusion because the results are highly consistent despite the fact that the MDS embedding is known to be non-unique and the embedding holds valid up to any arbitrary orthogonal transformations.

## Analysis of the top-3 group-wise topographic vectors

To derive a reliable measure of the whole-brain white matter topography, we systematically develop the novel methods for computing the group-wise topographic vectors. A total of 861 subjects are selected for the experiments presented in this section. The technical details and experiment settings are presented in the Methods section and Supplemental Information.

In this work, we are most interested in the topographic organisation present in the top-3 dimensions of the topographic vectors. This preference is based on the fact that the top-3 dimensions are significantly more representative than the remaining dimensions in terms of preservation of distance-metric geometry, as shown in **Figure 2** (B) and (C). It is also motivated by the widely-endorsed theory proposed in previous biological studies^15-17^ that the development of human brain white matter follows three principle chemotactic gradients established during the embryogenesis. The top-3 group-wise topographic vectors computed using our method for all the subjects in this study are shown in Figure 3 (A). Each topographic vector corresponds to a single streamline. Compared with the individual top-3 topographic vectors shown before in **Figure 1** (B) and here in Figure 3 (A), the group-wise topographic vectors form significantly more regular boundary and internal organisation. However, we are unable to clearly observe the internal organisation of the topographic vectors from the point cloud visualisation due to occlusion of points.

**Figure 3:**
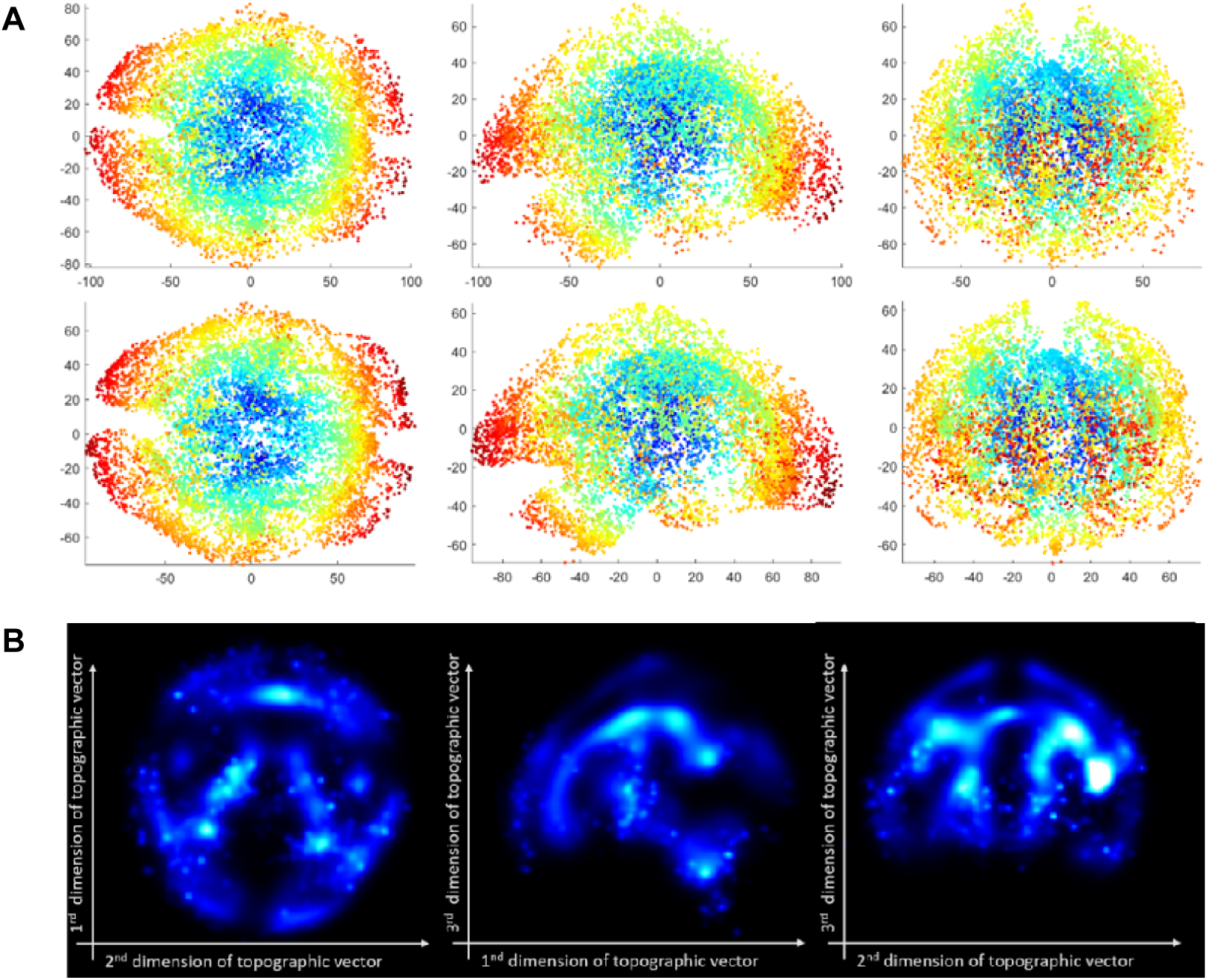
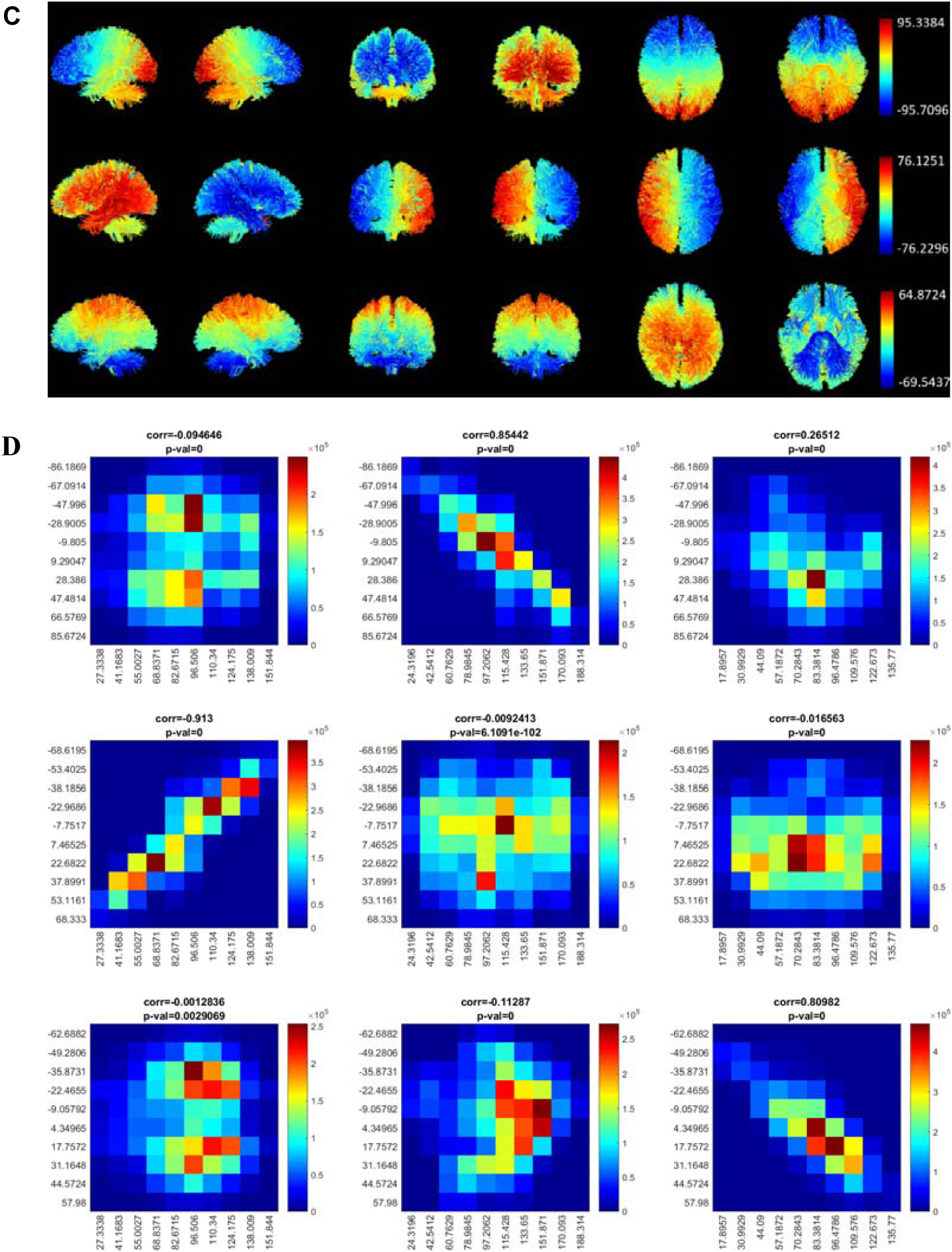
A) The point cloud view of the top-3 group-wise topographic vectors. B) The topographic vector density map view of the top-3 group-wise topographic vectors. C) The top-3 group-wise-topographic-vector-colourised whole-brain tractograms. D) Histograms of the topographic vectors against the brain anatomical coordinates of tractogram streamlines. The correlations and the p-values are displayed on top of each histogram.

For visualising the internal structure of topographic vectors, we also propose a density map visualisation of the topographic vectors, which we call the topographic vector density map (TVDM). The TVDM is computed from all of the aligned topographic vectors across different subjects using the adaptive multi-dimensional kernel density estimation^c 25^. The TVDM for all the subjects in our study is shown in Figure 3 (B). Similar to the point cloud view shown in **Figure *3*** (A), we can observe the brain shape of the enclosing boundary of the TVDM. From the sagittal view in the middle, we can also see a thin band of outer layer separating from the internal structures. Moreover, we find that several internal structures shown in the TVDM resemble the anatomical structures of the brain. For example, on the left of Figure 3 (B), we observe a few structures showing a remarkable resemblance to that of visual pathway and its occipital lobe projection. On the middle of Figure 3 (B), we observe the structures visually corresponding to the anatomy of the cingulate region and corticospinal pathways. On the right of Figure 3 (B), we observe the structures corresponding to the anatomy of the somatosensory pathways.

To visualise the association between the topographic vectors and the streamlines, we can colourise the streamlines using the values of each dimension of the topographic vectors. The colourised streamlines can be nicely drawn using QIT ^26^. Here we present the results for the top-3 dimensions of the topographic vectors in Figure 3 (C). The first to the last column of Figure 3 (C) shows different views of the topographic-vector-colourised tractograms. With this visualisation, we can observe smooth variations, a.k.a. the *gradients*, of the topographic values over the tractogram. Specifically, we can observe the gradients extending in the three anatomical directions: anterior-posterior, left-right, and superior-inferior. The relationship between the top-3 topographic vectors and the anatomical coordinates of the streamline points has been verified and precisely quantified in our study on individual topographic vectors shown in the previous section. The topographic vectors derived from the individual tractograms may suffer from various types of random errors, and they are unreliable. With the group-wise topographic vectors, we can derive a more reliable measure of the relationship. Similar to the individual study, we apply the Pearson linear correlation to each dimension of the top-3 group-wise topographic vectors and the dimensions of the streamline coordinates of the reference subject.

The 2D histograms of the association between the group-wise topographic vectors and streamline coordinates are shown in Figure 3. Strongly linear correlation between the topographic vectors and the respective anatomical coordinates of the streamline points can be ascertained based on both the histogram plots and the correlation values, echoing the statistical results for individual topographic vectors. The other weaker correlations are also consistent with the results for individual topographic vectors with stronger (closer to 0) *p*-values.

## Remaining dimensions of the topographic vectors

Although we are mainly focused on the top-3 dimensions of the topographic vectors, our method naturally applies to all the dimensions available in the MDS embedding. Anatomically meaningful patterns can also be seen in many of the remaining dimensions of the topographic vectors. Similar to Figure 3 (B), here we present the visualisation of the remaining 4^th^-10^th^ dimensions of the topographic vectors by colourising the streamlines with the corresponding topographic values in each dimension in Figure 4.

**Figure 4:**
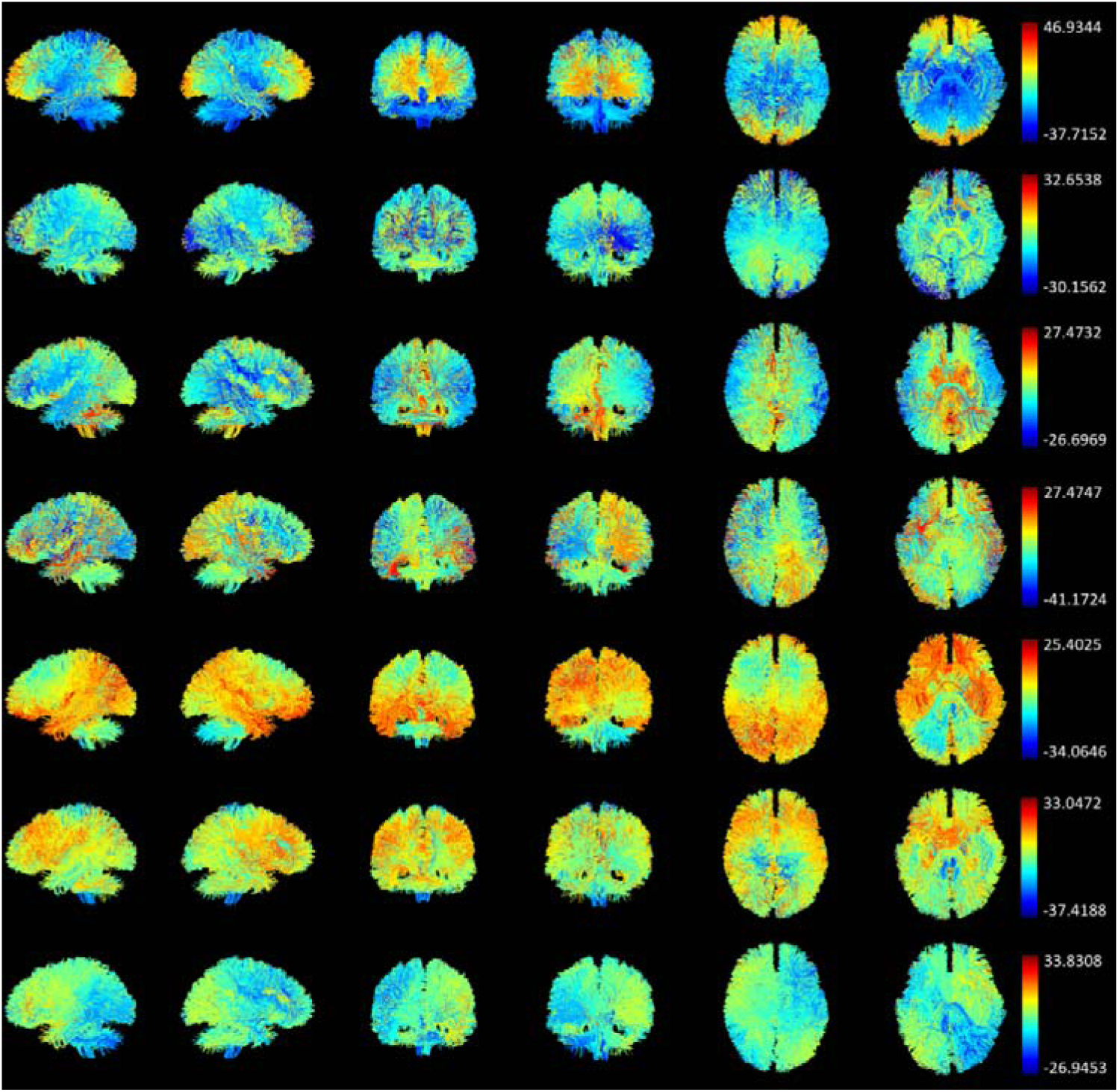
From the first row to the last row shows the whole-brain tractograms colourised by the 4^th^ – 10^th^ dimensions of the group-wise topographic vectors.

The 4^th^ dimension separates the motor streamlines passing through brainstem and cerebellum from all other streamlines and this pattern can be clearly observed; The 5^th^ dimension is quite complex, but the contrast in the occipital lobe, temporal and parietal lobe can still be seen from the sagittal views and coronal views; The 6^th^ dimension shows stronger contrast than the 5^th^ dimension, and the contrast differentiates the medial and lateral streamlines, as can be seen from the dorsal-axial view; we can also observe a difference between the left and right hemispheres from the posterior-coronal view; The 7^th^ dimension shows the most complex organisation, and we can barely observe a subtle segregation between the left and right hemispheres from the posterior-coronal view; The contrast patterns in the 8^th^ and 9^th^ dimensions are relatively clear and visually complementary in the sagittal and coronal views, while different highlights can be seen in the axial views; Specifically, the streamlines projecting to the superior frontal lope and cerebellum are highlighted in the 8^th^ dimension, and the streamlines projecting to the posterior parietal lope and brainstem are highlighted in the 9^th^ dimension; The contrast in the 10^th^ dimension is again visually clear in all views, a pattern of “X” shape is present: streamlines projecting from left-frontal to right-temporal and right-cerebellum (yellow-red), and from right-frontal to left-temporal and left-cerebellum (green-blue) are highlighted, although the contrast is modest.

Our analysis of the higher dimensions of the topographic vectors suggests that the higher dimensions of the topographic vectors encode the more subtle organisation of the white matter streamlines. It is more critical to connect patterns in the higher dimensions of the topographic vectors to the functions of the corresponding white matter tracts.

## Assessment of the reliability of our method

The validity of the findings reported in this work relies on the reliability of the methods that we developed for this study. In this section, we present our experimental results in this regard. A total of 861 subjects are selected for the experiments presented here. The details on the experiment setting are presented in the Methods section.

### Test-retest reliability of the individual topographic vectors

We first report the reproducibility of the individual embedding for the same subject. In this experiment, we compare the individual topographic vectors of the two whole-brain tractograms computed by two separate runs of the tractography of 100 different HCP subjects. The topographic vectors of the second-run tractography are aligned and matched to those of the first-run tractogram based on the topographic vector alignment and matching algorithm developed for the group-wise topographic vector analysis. Based on the guideline of the statistical testing for evaluating reproducibility ^27^, for assessing test-retest reproducibility we compute the 2-way mixed-effects model of the intra-class correlation coefficient, a.k.a. ICC(3,k), and we chose the absolute agreement to be the definition. We compute the ICC values and the lower and upper bounds for *α* = 0.05. The boxplots, mean values, lower and upper bounds of the ICC are shown in Figure 5 (A) and (B).

**Figure 5:**
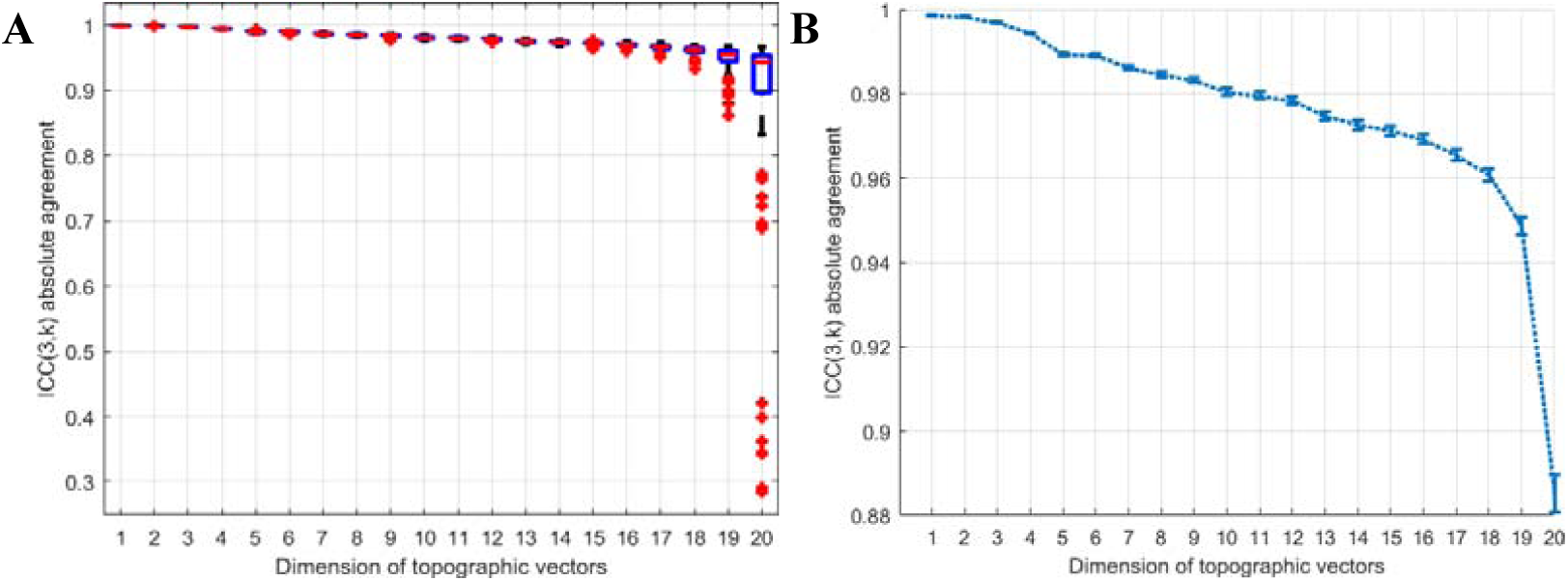
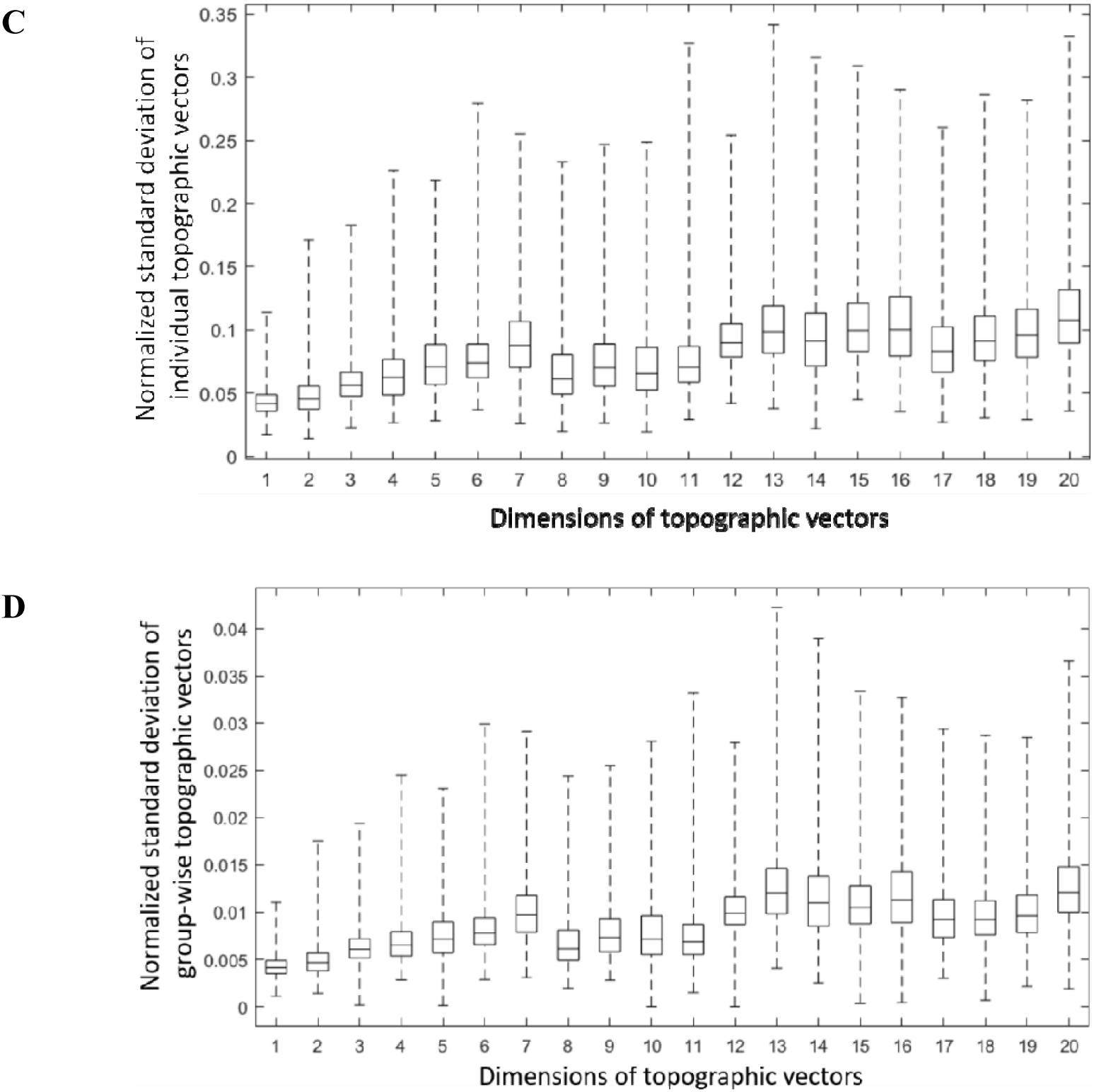
Reproducibility of individual topographic vectors. A) Boxplot of the ICC values for each dimension of the topographic vectors; B) The means of the ICC values are shown as a dashed line and the lower and upper bounds of the ICC value with significance level α=0.05 are shown by the spreads of the bars at each point of the mean ICC values. C) Boxplot of the standard deviation of aligned individual topographic vectors. D) Boxplot of the standard error of bootstrapping group-wise topographic vectors.

The ICC values indicate excellent reproducibility of the topographic vectors in general. However, we observe a sharp decline at the 19^th^ dimensions of the topographic vectors. More outliers of the ICC values can be seen in the 19^th^ and 20^th^ dimensions. From the plots of the mean and standard deviation of the relative differences, we observe a similar decline of reproducibility in dimensions 19 and 20. In addition, we observe a margin between the 4^th^ and 5^th^ dimensions in the ICC values.

### Reliability of group-wise brain streamline topography analysis

We are most concerned about the question: will the group-wise topographic vectors be the same for different samples of the same population? We address this concern by using bootstrapping. In this experiment, we resample the whole study dataset to obtain 100 individual subjects with replacement for 1000 times. This procedure gives us 1000 different subsample sets of the study set, each of which contains 100 individuals. We then compute the group-wise topographic vectors for the 1000 subsample sets. Afterwards, we compute normalized standard deviations of the topographic vectors up to the first 20 dimensions of the topographic vectors across the 1000 subsample sets, or the standard errors. The results are shown in the boxplot of Figure 5 (D). We also present the normalized standard deviations of the aligned individual topographic vectors in Figure 5 (C) for comparison ^d^. The group-wise topographic vectors are significantly more reliable than the individual topographic vectors. The normalized standard deviation of each dimension of the topographic vectors is defined as the standard deviation of the values of the topographic vectors divided by the maximum magnitude of the values of all the samples in each dimension.

## Discussions

### Methodological

Note that the individual streamlines computed by the tractography algorithm do not represent individual axonal connections. Instead, each streamline represents a small bundle of axons at mesoscale aligning with the magnetic field gradient directions measured by diffusion MRI. Accordingly, we would view our measure as a mesoscale measure of the global topographic organisation of the whole brain streamlines.

Our study may be considered an application of the dMRI tractography. Albeit its uniqueness for *in vivo* and non-invasive measurement of brain anatomical connectivity, a well-known criticism towards tractography and its applications is that presently the accuracy of tractography with dMRI may still be limited ^28-31^. The limited accuracy of the tractography has been attributed to the bias of tractography that it terminates preferentially on gyral crowns, rather than the banks of sulci ^29,31^. It may be also attributed to the nonunique interpretations of the streamline topology within single voxel from the measured diffusion MRI signals at the voxel, causing a significant amount of false-positive connectivities^30^. In the past decades, tremendous efforts have been made to improving the quality of diffusion MR imaging^19,30,32-36^ and tractography algorithm^37-39^. The bias in the tractography algorithms and the ambiguity in the resulting streamline topology may be considerably further reduced or even resolved to a degree of satisfaction with the advances in diffusion MR imaging and computing technologies.

In this work, we have thoroughly examined the reproducibility of the proposed method for the HCP data, and the results are satisfactory. However, the multisite reproducibility of the dMRI data and the resultant analysis is still an open challenge ^40,41^. The proposed method may also be used for the task of multi-site harmonization since the tractograms are all mapped to the MDS embedding space in which the white matter tracts can be more comparable than in the streamline space.

Previously, spectral embedding has been proposed for streamline embedding, and the embedding vectors are used for tractogram clustering ^42^. In spectral embedding, only the proximity relationship, in the form of normalized affinity, between close-by streamlines is considered, and the global organisations are neglected due to their close-to-zero affinity values of far apart streamlines. On the contrary, all pairwise distances are to be preserved in the embedding vectors in MDS. Accordingly, we consider the MDS embedding as a more suitable method for measuring the whole brain streamline topography. MDS has also been used for visualising individual tractograms and at most 3 dimensions has been considered ^43^. However, the MDS was not immediately applicable to group-wise study due to the computational intractability, although the group-wise study with high reproducibility is crucial to a reliable measurement. Furthermore, the anatomical interpretations of the embedding vectors were not elucidated in the previous works.

### Biological

From the histograms presented in Figure 2 (D), we also observe that the strongly correlated dimension pairs of the streamline coordinates and topographic vectors are very stable without any alignment. The highly consistent individual MDS embedding results across different subjects suggest that the eigenvectors and eigenvalues obtained by eigendecomposition of the Gram matrices of different subjects are unusually stable. According to the Rayleigh quotient and min-max theorem, the magnitudes of the eigenvalues characterize the significance of the corresponding subspaces of the original distance matrix. Hence, we can conclude that the top-3 most representative subspaces of the metric geometry of the streamlines are highly consistent. If we view the strongly and linearly correlations between the dimension pairs of the streamline coordinates and the topographic vectors as a law of whole-brain white matter wiring, the consistency of this result would imply that different brains obey the same law of whole-brain white matter organisation.

In our study of the group-wise topographic vectors, we observe a locally smooth and regular shape formed by point cloud of the group-wise topographic vectors. The local smoothness and regularity of the topographic vectors in the embedding space implicate the local topographic regularity of the streamlines. We also observe that the outer layer of the topographic vector point cloud resembles the shape of the brain. This finding is not surprising since the surface of the white matter is filled with short association streamlines. When representing these short streamlines as 3D vectors using MDS, the geometrical organisation of these 3D points ought to agree with the actual positioning of the corresponding streamlines in the tractograms. Since the short association streamlines are all located near the white matter surface, the corresponding topographic vectors will also bear the same geometry, forming the shape of the white matter surface. In addition, from the TVDM drawn based on the topographic vectors of all subjects in this study, we observe that the shapes of the internal structures of the topographic vectors are also regular. This observation can also be viewed as evidence of regularity in the global topography of the whole brain streamlines in the brains of young adult population.

Our method applies to all dimensions of the topographic vectors. Anatomically meaningful patterns have been observed in many of the dimensions of the topographic vectors. It is also possible to further study the relationships between the higher dimensions of the topographic vectors and the anatomical coordinates of the whole brain streamline streamlines. However, due to the lack of biological interpretations, we decide to include these results in the Supplementary Information.

## Methods

### Dataset and preprocessing

We adopted the 900 Subjects Data Release of the original young adult HCP dataset in our study. We selected the subject with dMRI scans of a total of 288 gradient directions for 3 b values, resulting in 861 subjects in total. The streamline orientation density (FOD) maps for all the dMRI data are reconstructed using the method proposed in ^44^. The order of spherical harmonics is set to 12. Afterwards, a whole-brain tractogram of 10,000 streamlines is generated for each subject by using the anatomically constrained tractography implemented in MRtrix3 with the default parameter setting^37,45^. This number of streamlines is chosen based on the common practice of connectome research and the tolerability of the computational cost.

For the group-wise study, we warp all tractograms to a common space. First, a track-density image (TDI) ^24^is generated for each tractogram and then a nonrigid warp field is generated by registering all TDI images to a reference TDI image using SyN implemented in ANTs^46^. Subject 100307 is selected as the reference for TDI. All tractograms are subsequently warped to the reference space by using the generated warp field, such that the individual differences and noises are removed as much as possible for producing a reliable group-wise measure. In our experiment, we adopt the one-sided Hausdorff distance *d*_*H*_ (*X,Y*) = max_*y∈Y*_ min_*x∈X*_ *d*(*x,y*) for measuring the distances from the streamline *X* to streamline *Y*. We symmetrize the one-sided Hausdorff distance by *d* (*X,Y*)= min (*d*_*H*_ (*X,Y*), *d*_*H*_ (*Y,X*)). The standard Hausdorff distance *d* (*X,Y*) = max (*d*_*H*_ (*X,Y*), *d*_*H*_ (*Y,X*)) is a common choice in white matter tract analysis. Our alternative formulation differs slightly from the standard Hausdorff distance in that the short streamlines are considered to be part of the close-by long ones. This reformulation is proposed based on the connectional anatomy of brain white matter pathways, and usually short tracts in the tractograms are considered as false positives of tractography^47^. This choice of distance measure is by no means unique^48^. The proper selection of the distance measure may be based on the function that the anatomical connectivity, in the form of streamlines, serves, which is out of the scope of this work. All the computations were carried out with the distributed parallel-computing infrastructure of the LONI server grid ^49^.

### Individual streamline topography analysis via MDS embedding

Our model of white matter topography analysis is based on the notion of multidimensional scaling (MDS), and the MDS maps each curve to a vector in a vector space of fixed dimensions. Since the MDS mapping is isomorphic and isometric for Euclidean geometry, the resultant finite-dimensional embedding space and the streamline curve space are equivalent in their metric geometry. Furthermore, each dimension of the embedding vectors encodes a subspace of the original streamline space. Because of these fine properties of the MDS embedding, we consider the resultant embedding vectors as the measure of the topography of the whole brain streamlines. In addition, we have shown that it is straightforward to interpret this measure to the known brain anatomy.

The MDS embedding of an individual streamline can be computed straightforwardly with the following steps ^22^: a) compute the gram matrix from the full pairwise distance matrix; b) perform eigendecomposition on the gram matrix; c) compute the embedding vectors using the positive eigenvalues and the associated eigenvectors. The detailed formulations and theoretical analysis are included in the Supplementary Information.

### Group-wise brain white matter topography analysis

To derive a reliable measure of whole-brain white matter topography, we propose to extend the individual topographic vectors to the group-level. The major problem in the way of achieving it is that the whole-brain tractograms and the associated topographic vectors are not immediately comparable across different subjects due to individual differences and geometrical complexity of the streamline curves in the whole-brain tractograms.

To remove the individual differences in the individual tractograms, we have warped the individual whole-brain tractograms to a standard anatomical space using volumetric registration in the preprocessing step, and the remaining cross-subject inconsistency in the tractography after the registration is considered a result of random errors. The individual topographic vectors from the aligned tractograms are more comparable. However, a known issue with the MDS embedding that we use for computing the topographic vectors is that the result is non-unique. Theoretically speaking, there are infinitely many plausible solutions to the MDS given a distance matrix. Therefore, the topographic vectors across different tractograms have also to be aligned. Intuitively, the alignment may be achieved by computing the MDS for the entire set of all the tractograms composed of all individual ones from the study at once. However, the other technical challenge we face is that it is generally intractable to apply MDS to this overwhelmingly massive tractogram since the full pairwise distance matrix of it is both prohibitive to calculate, store and to use in the MDS. Alternatively, we propose to align and match the individual embedding vectors across different subjects. Afterwards, the group-wise topographic vectors can then be computed by averaging the matched topographic vectors across all subjects in the study. The technical details are included in the Supplementary Information.

### A fast streamline k-NN algorithm

Our group-wise streamline-wise topography analysis requires the k-nearest-neighbour (k-NN) streamline distances across different streamline bundles. However, it is often intractable to compute the *k-NN* streamline distances for a massive set of tractograms^48,50^. Based on the observation that the streamline-wise *k-NN* distance is strongly related to the point-wise *K*-NN distance, we propose a fast streamline *k*-NN algorithm. This algorithm constitutes the core component of our group-wise topography analysis framework due to its affordable computational cost.

Conventionally, to compute the *k-NN* streamlines we will need to calculate all pairwise distances between streamlines, and the pairwise streamline distances are generally calculated by resorting to all pairs of points on the streamline pairs, which is computationally expensive. Alternatively, we propose a *k*-NN-likeness measure for streamlines. Based on this measure, we can define the approximate streamline *k-*NN to a given streamline. The advantage of this approximate streamline *k*-NN definition over the exhaustive streamline *k*-NN is that we only need to compute the streamline *k*-NN-likeness measure for a limited subset of streamline pairs, and it is also trivial to compute this measure. The complete fast streamline *k*-NN algorithm and more theoretical discussions, as well as the experimental results, are given in the Supplementary Information.

## Code availability

Source code of the proposed methods is available online at https://www.nitrc.org/projects/connectopytool/, along with the code for reproducing the results presented herein.

## Supplementary Information

is available in the online version of the paper.

## Acknowledgements

This work was in part supported by the National Institute of Health (NIH) under Grant RF1AG056573, R01EB022744, U01EY025864, U01AG051218, P41 EB015922, P50AG05142. Data used in this paper were provided by the Human Connectome Project, WU-Minn Consortium (Principal Investigators: David Van Essen and Kamil Ugurbil; 1U54MH091657) funded by the 16 NIH Institutes and Centers that support the NIH Blueprint for Neuroscience Research; and by the McDonnell Center for Systems Neuroscience at Washington University.

## Author contribution

J.W. developed the new methods, J.W. and Y.S. designed the experiments, J.W. performed the analysis, J.W. and Y.S. co-wrote the paper, Y.S. supervised the project.

## Author Information

The authors declare no competing financial interests.

## Supplementary information

### S1 Introduction

#### Streamline *k*-NN

Computing the k-nearest-neighbour (*k*-NN) distance for the streamlines in the massive tractograms from a population of subjects is often inevitable in large-scale white matter analysis using tractography based on dMRI ^48,51-56^. However, this procedure is known to be computationally demanding, and it can be intractable for large datasets. A common practice to alleviate the problem is to re-sample the streamlines of varied lengths and topology to the same number of points, such that the distances can be calculated by taking the vector norm of the point-wise distances along the streamlines. The rationale of this method lies in the fact that the re-sampled streamlines still represent the original streamlines to a certain degree, and the normed point-wise distance indeed captures the relative closeness of the streamlines in comparison. A limitation of this method is that the order of the points on nearby streamlines can be arbitrary. Hence, it is required to re-order the points on the streamlines before taking the norm, which can be inconvenient and error-prone. Moreover, resampling potentially causes significant ambiguity in streamline comparison. In **Error! Reference source not found.**, we illustrate a straightforward example of such ambiguity in streamline comparison caused by streamline re-sampling. It shows that the apparent representations of the streamlines, shown as dots in a two-dimensional virtual plane, form an equilateral triangle, while from an anatomical point of view the streamlines would be better to lie on a straight line in the 2D virtual plane with the blue dot in the middle.

**Supplementary Figure 1.**
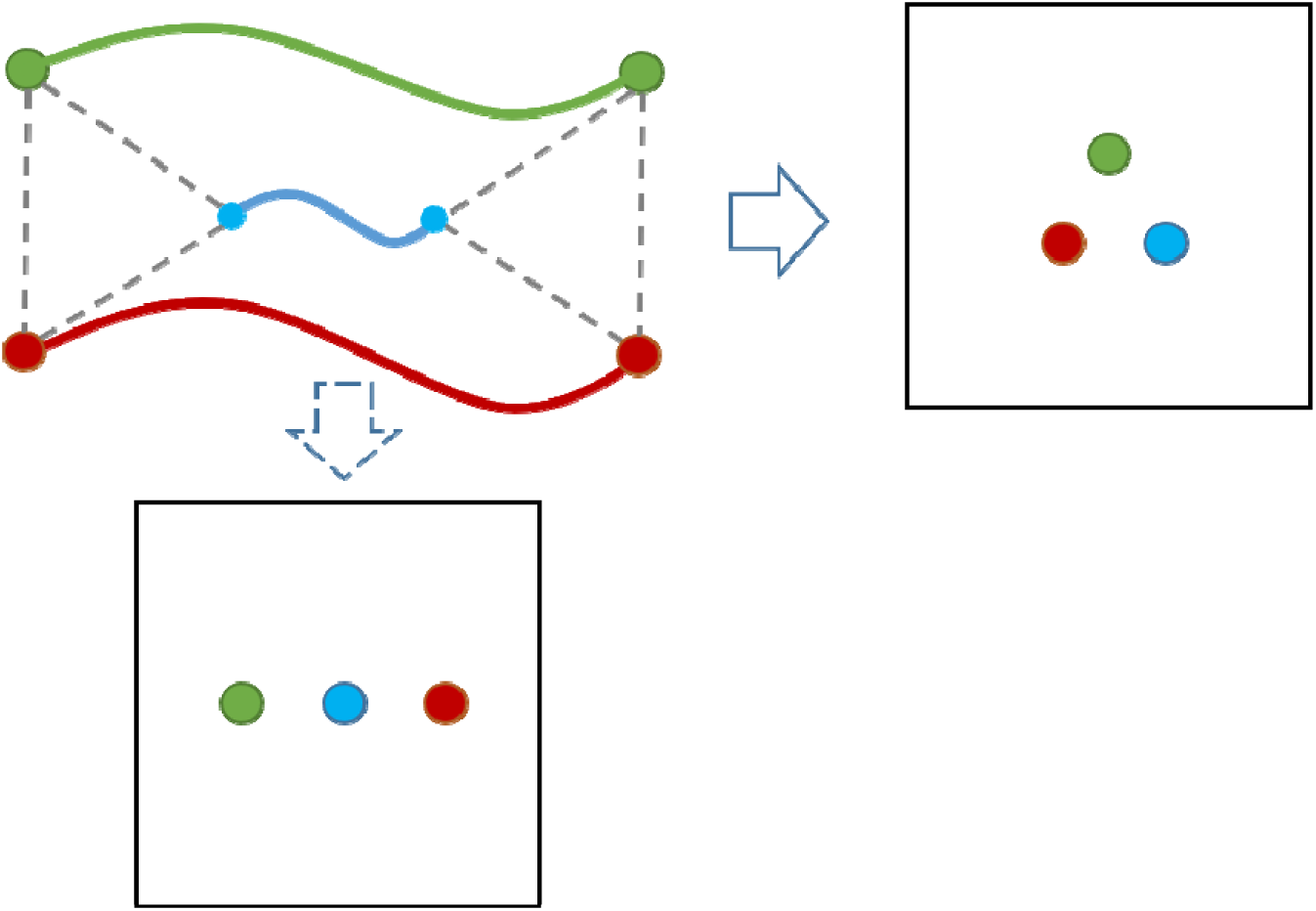
Ambiguity in streamline distance due to re-sampling. Green, blue, and red lines in the top-right figure are three streamline curves. The extreme points on the lines are the re-sampled points. The dash lines represent the point-wise distances between the re-sampled points on the different streamlines. The green, blue and red dots in the two other boxes are 2D isometric point visualization of the streamlines in the same colours.

As the main contribution of the method development, we propose a fast streamline *k*-NN algorithm, which is the first of this type to the best of our knowledge. Our fast streamline *k*-NN algorithm is motivated by the observation that the streamline-wise *k*-NN distance is strongly related to point-wise *K*-NN distance. In the remainder of this paper, we use *K*-NN to denote point-wise *K*-NN and *k*-NN to denote streamline-wise *k*-NN. In this work, we first establish the theoretical relationship between point-wise *K*-NN and streamline-wise *k*-NN. Accordingly, we propose a streamline *k*-NN-likeliness measure which can be computed efficiently with point-wise *K*-NN algorithms, such as the celebrated k-d tree algorithm ^57^, and this gives rise to our fast streamline *k*-NN.

#### Whole-brain white matter topography analysis

As the second contribution of the method development, we propose an anatomically meaningful streamline-wise measure of the white matter topography. In addition, we extend our method for individual white matter topography analysis to scalable group-wise analysis using our proposed fast streamline *k*-NN algorithm.

Our model of white matter topography analysis is based on MDS embedding ^21,22^ of the streamline curves. The MDS maps each streamline to a vector in a finite-dimensional vector space. Since the mapping is isomorphic and nearly-isometric, we consider that the resultant finite dimensional embedding space and the streamline curve space are equivalent in their geometrical construct, and the topography of the white matter tracts are characterized by the statistical and geometrical analysis of the embedding vectors. Although streamline embedding has been applied to white matter tract segmentation and visualization ^42,43,58^, this is the first time that it is used as a measure of white matter topography and it is also the first time that the anatomical meaning of the MDS embedding of the white matter tracts is elucidated and quantified to the best of our knowledge. Furthermore, we extend the streamline embedding of individual subjects to group-wise embedding to derive a statistically reliable measure of the white matter topography. The major problem in the way of achieving it is that the whole-brain tractograms and the associated topographic vectors are not immediately comparable across different subjects due to individual differences and geometrical complexity of the streamline curves in the whole-brain tractograms.

To remove the individual differences in the individual tractograms, we have warped the individual whole-brain tractograms to a common anatomical space using volumetric registration in the preprocessing step, and the remaining cross-subject inconsistency in the tractography after the registration is considered a result of random errors. The individual topographic vectors from the aligned tractograms are more comparable. However, a known issue with the MDS embedding is that its solution is non-unique. Theoretically speaking, there are infinitely many plausible solutions to the MDS given a distance matrix. Therefore, the topographic vectors across different tractograms have also to be aligned. Intuitively, the alignment may be achieved by computing the MDS for the entire set of all the tractograms composed of all individual ones from the study at once. However, the other technical challenge we face is that it is generally intractable to apply MDS to this overwhelmingly massive tractogram since the full pairwise distance matrix of it is both prohibitive to calculate, store and to use in the MDS. Alternatively, we propose to align and match the individual embedding vectors across different subjects. Afterwards, the group-wise topographic vectors can then be computed by averaging the matched topographic vectors across all subjects in the study.

#### Whole-brain streamline matching by topographic vector alignment and matching

As a useful by-product of our work, the topographic vector alignment and matching can be viewed as a novel scalable method for non-rigid whole-brain tractogram matching. Identifying corresponding streamlines across different massive tractograms is a critical task for group-wise white matter analysis. However, due to the well-known challenges, such as large variability of the streamline curves from tractography, ambiguity in streamline-wise comparison as well as the computational intractability, the research on non-rigid whole-brain white matter tractogram matching is still sparse. In most of the existing methods for direct streamline-wise registration, only linear, i.e. rigid or affine, transformations are considered to model the variability of the streamlines^50^. This assumption is mostly valid for longitudinal studies. However, it generally does not hold for different subjects in cross-sectional studies. More recently, the tractogram matching problem has been cast as a graph matching problem^59^. Although this idea is mathematically sound, it deals with the whole brain tractograms smaller than the ones adopted in common whole-brain structural connectivity analysis^60^. The limited scalability of their method is perhaps mainly due to the high computational complexity of graph matching algorithms, and the need for exhaustive pairwise streamline distance computation as a prerequisite. Hence, it is unscalable to large-scale analysis. Alternatively, a nonrigid volumetric registration may be performed, and the linear streamline-wise matching can be done subsequently. A drawback of this method is that it is generally assumed that no topological changes occur in the images to be registered, such that a voxel in one image can always be matched to a voxel in the other image. Since the topology of white matter tracts varies considerably across different subjects, it is arguable if the volumetric registration is applicable to streamline matching.

A preliminary version of this work was presented in^61^. In this work, we substantially extend the previous work by fundamentally improving the group-wise topographic vectors via a novel topographic vector alignment and matching framework, and the latter by itself is novel as a method for non-rigid whole-brain tractogram matching.

#### S2 The Fast streamline *k*-NN algorithm

Conventionally, to compute the streamline *k*-NN we will need to compute all pairwise distances between streamlines, and the pairwise distances are generally computed by resorting to all pairs of points on the streamlines, which is computationally expensive.

##### The theoretical relationship between the *K*-NN point distance and the *k*-NN streamline distance

Without loss of generality, in the theoretical discussions presented below, we treat a streamline curve as a point set, and we also consider a set of streamlines, or tractogram, as a collection of point sets. Our main idea of the fast approximation of the streamline *k*-NN distance lies in the following theorem.

###### Theorem 1.

*Suppose we are given a collection of point sets, denoted as* **Γ**, *and two point sets* X, Y *belonging to* **Γ**, i.e., *X, Y* ∈ **Γ** *and suppose* 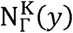 *is the set of the K-NN points of a point y* ∈ *Y within* **Γ** *and the point-wise K-NN is defined by point-wise distance d*(·, ·), *additionally if* 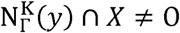, *then*

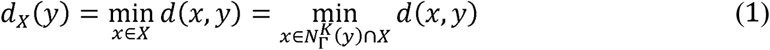

This theorem shows that we do not require the entire set, or streamlines, *X* to compute *d*_*X*_(*y*), but we would only require the pointwise *K*-NN computed beforehand, which lays the theoretical basis of our method. The proof is deferred to the appendix.

According to Theorem 1, if 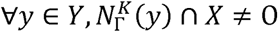, we can rewrite a common streamline distance as follows:

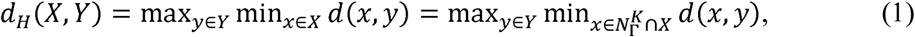

which is also known as the one-sided Hausdorff distance.

Besides, we observe that 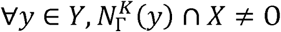 implies that *X* and *Y* are close-by. This can be described formally as

###### Proposition 1.

*Suppose* 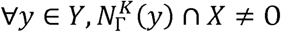,

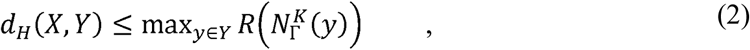

*Where* 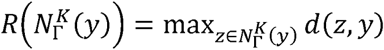 *may be called the radius of* 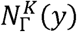.

Proposition 1 can be proven by upper bounding *d*_*H*_(*X,Y*) with the definition of *d*_*H*_.

##### Fast streamline *k*-NN approximation

Motivated by the observations presented in the last subsection, we propose to construct the streamline *k*-NN efficiently, and we can naturally relate the size of 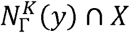. To the closeness of streamlines, and we propose a streamline *k*-NN-likeness measure based on the size of 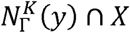, which we will later show can be computed efficiently, as follows:

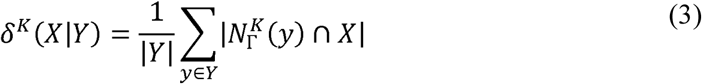

where |·|is the cardinality, or size, of a set. *X* with larger *δ*^*k*^(X|Y) is considered closer to *Y*, which gives rise to our definition of the key notion of this work:

###### Definition 1 (Approximate streamline k-NN).

*We define the approximate streamline k-NN in* Γ *for streamline Y as the set* 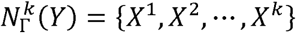, *if* ∃*K such that δ*^*k*^(X|Y) ≥ *δ*^*k*^(X′|Y)*for* 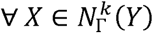 *and* 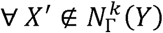

The *k* is usually much smaller than the *K* value in the point-wise *K*-NN.

In the following, we show the main point of this work that Eq. (3) can be computed efficiently without resorting to all pairwise streamline distances based on the following observations.

###### Definition 2 (Point-to-tract mapping).

*For streamline bundle T* = {*t*^1^, *t*^2^, …*t*^*N*^}, *where each streamline is defined by a point set as* 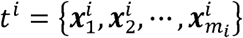, *if we denote the set of all points on all streamline streamlines as* 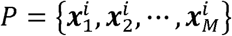, *the injection mapping* PT:*i* ∈ {∀*i*′|*x*_*i*′_ ∈ *P*} ↦ *j* ∈ {∀*j*′|*t*_*j*′_ ∈ *T*}*is called a* ***point-to-tract*** *mapping*. The tractograms are stored as a point set in benchmark file formats, such as the `trk` and `tck` files, and this mapping is inherently provided with the tract files.

###### Proposition 2 (Indicator of tract neighbours).

*Suppose* 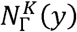 *is a set of point indices and* 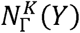 *denotes the union of the indices of tracts containing points belonging to the K*-NN *of certain y* ∈ *Y*, i.e. 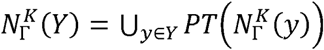.*We have*

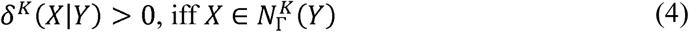

This proposition can be proven by definition of the point-to-tract mapping *PT*, i.e. Definition 2, definition of 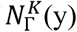 and definition of *δ*^*K*(*X*|*Y*)^. It shows us that we can easily identify the union set 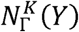 and compute Eq. (3) for this set only. Moreover, we can see that it is also straightforward to compute 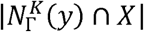 for Eq. (3), since

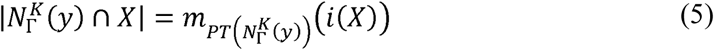

where *i*(*X*) is the index of *X* and *m*_*A*_(*x*) is the multiplicity of *x* in the multiset A^e^.

**Supplementary Figure 2:**
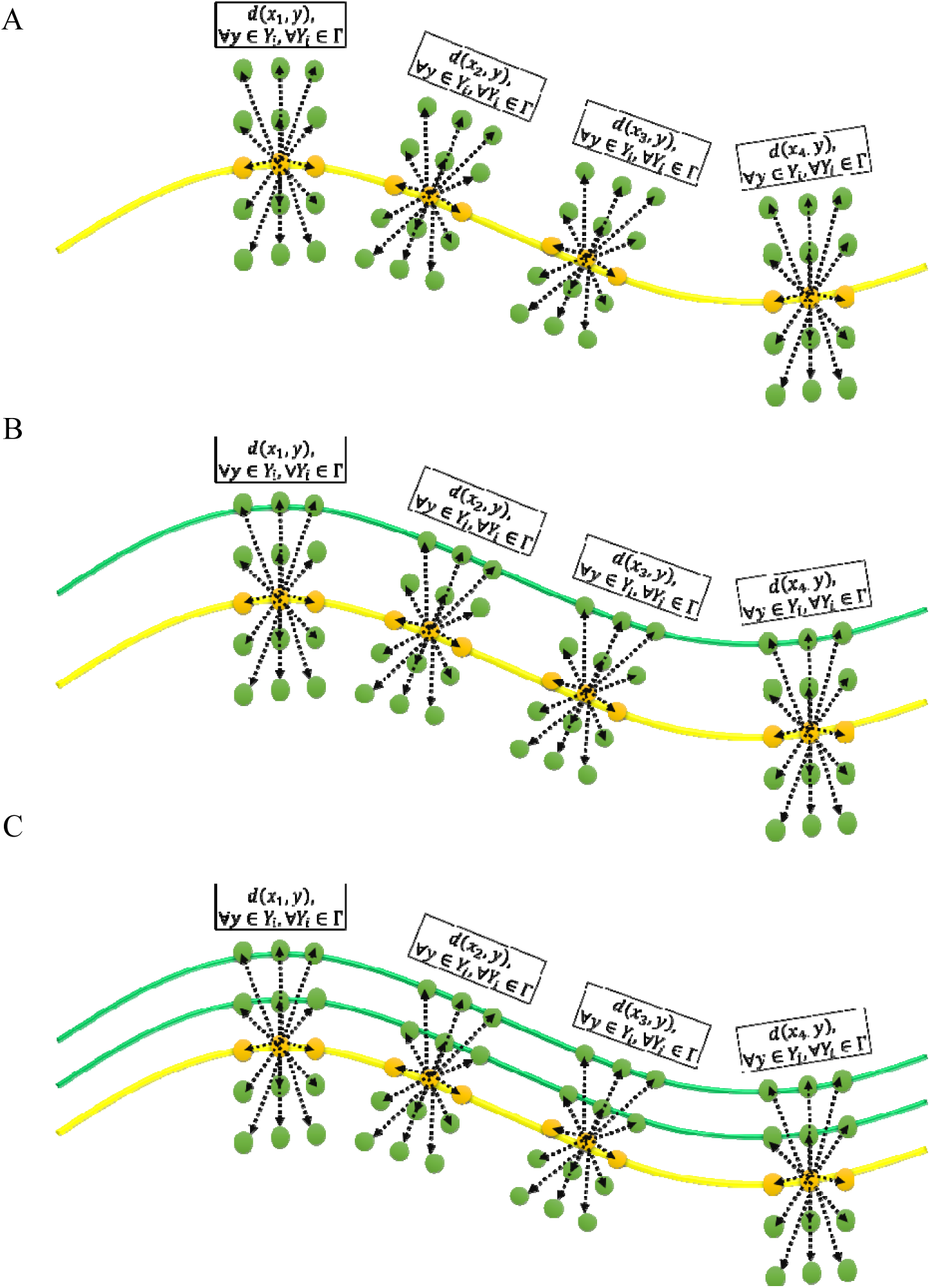
Illustration of the fast streamline *k*-NN algorithm: A) Compute point-wise *K*-NN for each point on a streamline; B) Identify the points within the point-wise *K*-NN on other streamlines; C) Sort the streamlines based on the number of points of the point-wise *K*-NN on the streamlines.

We illustrate the idea of the fast streamline *k*-NN algorithm in **Supplementary Figure 2**. In a nutshell, instead of searching for the neighbouring streamlines based on the streamline distances, we identify the neighbouring streamlines based on the easy-to-compute measure of likeliness of neighbourhood, i.e. Eq. (3). Finally, we summarise our fast streamline *k*-NN algorithm in Algorithm 1.

###### Algorithm 1: Fast streamline *k*-NN algorithm

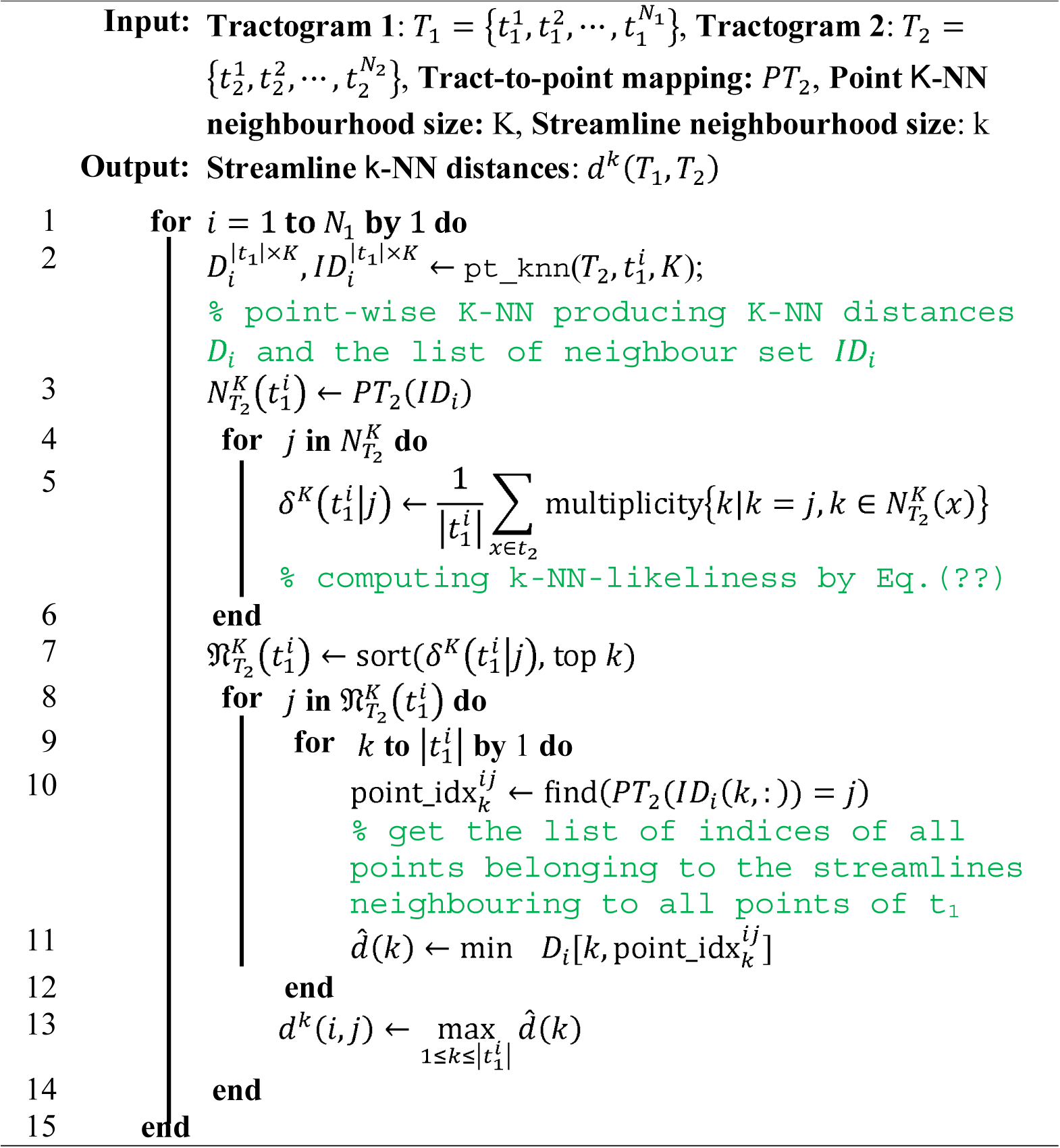

##### Reliability of the fast streamline k-NN algorithm

Here we present the experimental results for assessing the performance of our fast streamline *k*-NN algorithm. In our experiment, we adopt the one-sided Hausdorff distances and omit the symmetrisation for simplicity. The point-wise neighbourhood sizes in our method are K=100, 200, …, 1000. The streamline-wise neighbourhood size for our method is always set to *k*_1_=10, and the neighbourhood sizes for the exhaustive streamline distance computation are *k*_2_=10, 20, …, 100. We first evaluate our method in terms of the number of overlapped streamline sets in the tract neighbourhood from both methods. The results are shown in **Supplementary Figure 3**. From the histograms, we observe that the neighbourhood overlap rate increases w.r.t. both *k*_*2*_ and *K*. Since in practice the streamline-wise distances can be recomputed upon determining the approximate *k*-NN. It is possible to accurately approximate streamline *k*-NN with a large *k*_2_ or K.

**Supplementary Figure 3:**
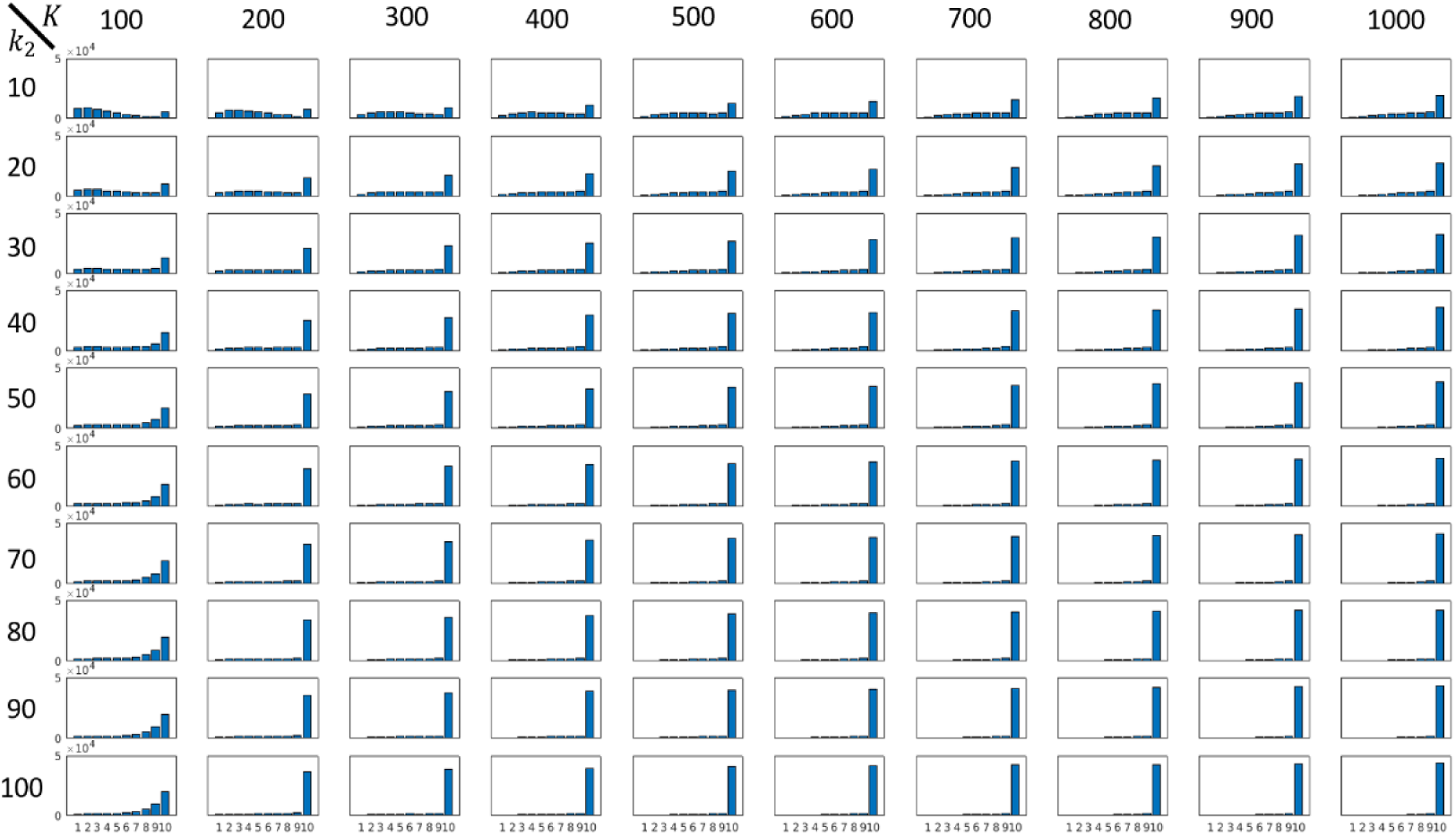
Histograms of neighbourhood overlap between our method and ground truth. The *x*-axis is the *k*-NN overlap, ranging from 1 to 10.

Furthermore, we study the relationship between the computational cost and the accuracy of our method in different parameter settings. First, we calculate the mean neighbourhood size overlap, **mean**(*k*\*), for different *K* values, and we also calculate the speed-up ratio of our method, denoted by *r*, which is defined as the computation time of exhaustive streamline distance computation divided by that of our method. The results are shown in Supplementary Figure 4 (A). The baseline computational cost for the exhaustive streamline distance computation is 15826.47 sec in MATLAB on a laptop PC with Intel i7-6820HQ CPU and 32GB ram. On the contrary, our method requires only couples of minutes to achieve 7 or 8 neighbourhood overlap. To further quantify the performance, we fit the scatter points of the mean neighbourhood size overlap *m***=mean**(*k*\*) and speed-up ratio *r* to two curves against *k*_2_ and K using least-absolute-residuals (LAR) robust fitting, as shown in **Supplementary Figure** 4 (B). Note that the speed-up ratio is independent of *k*_2_, since all pairwise distances are computed for any *k*_2_. The results are summarized in **Supplementary Table *1***.

In addition, we present the results for inter-subject fast streamline *k*-NN in **Supplementary Figure 5**. In this experiment, we compare the results of the fast streamline *k*-NN with the ground-truth streamline *k*-NN for 100 subjects from the HCP dataset. From the results, we observe that the overlaps between the top-20 *k*-NNs of our method and top-20 ground-truth *k*-NNs are generally high. The overlaps are larger for the 50K tractogram than the 10K tractogram. This means that our fast streamline *k*-NN is more effective for denser tractograms.

**Supplementary Figure 4:**
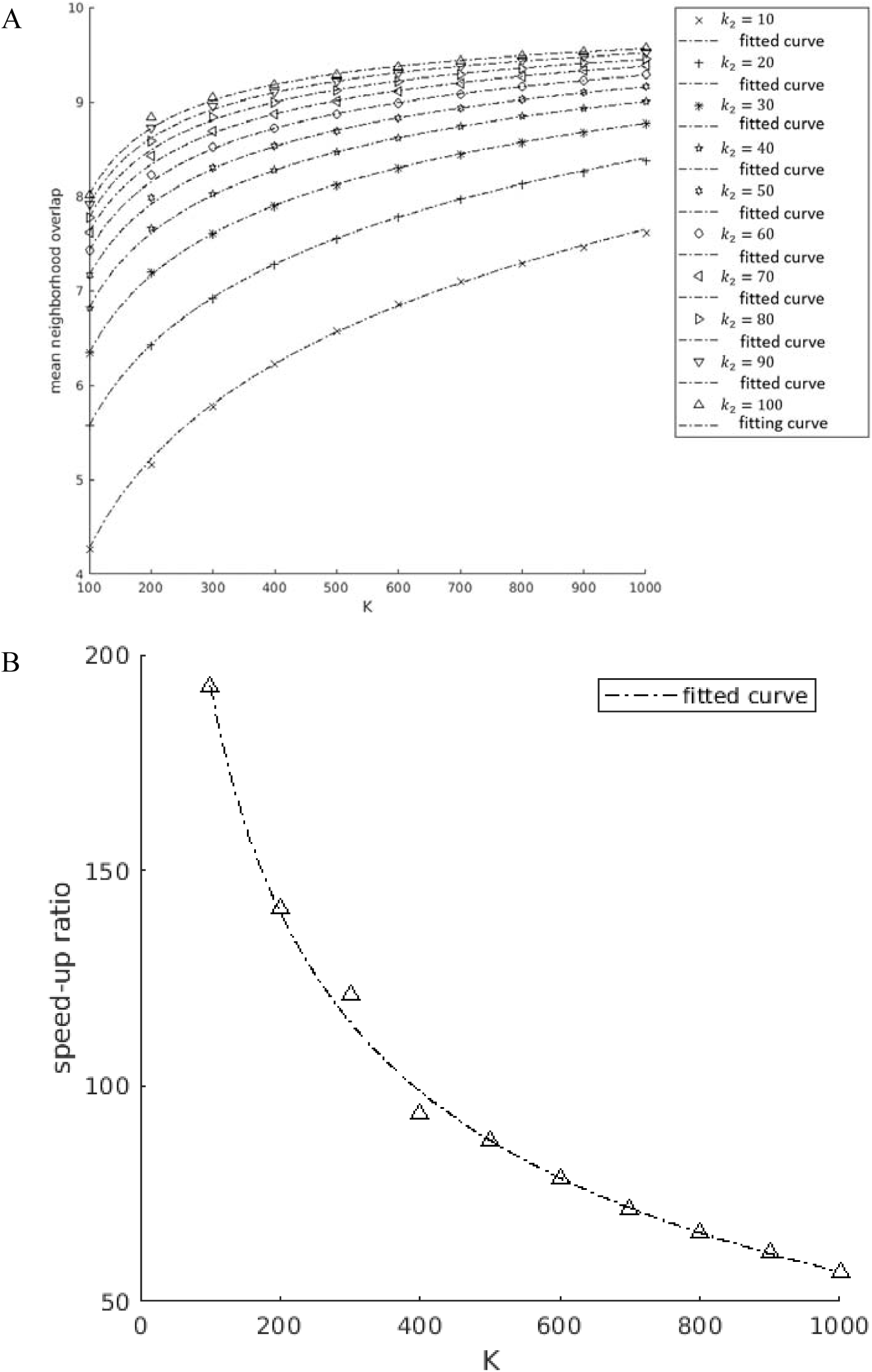
Performance curves of the fast streamline *k*-NN algorithm. (A) The neighbourhood overlap of our method. (B) The computational speed-up ratio of our method against the exhaustive pairwise streamline distance computation. The dashed curves are LAR robust fitting of the scatter points.

**Supplementary Table 1:**
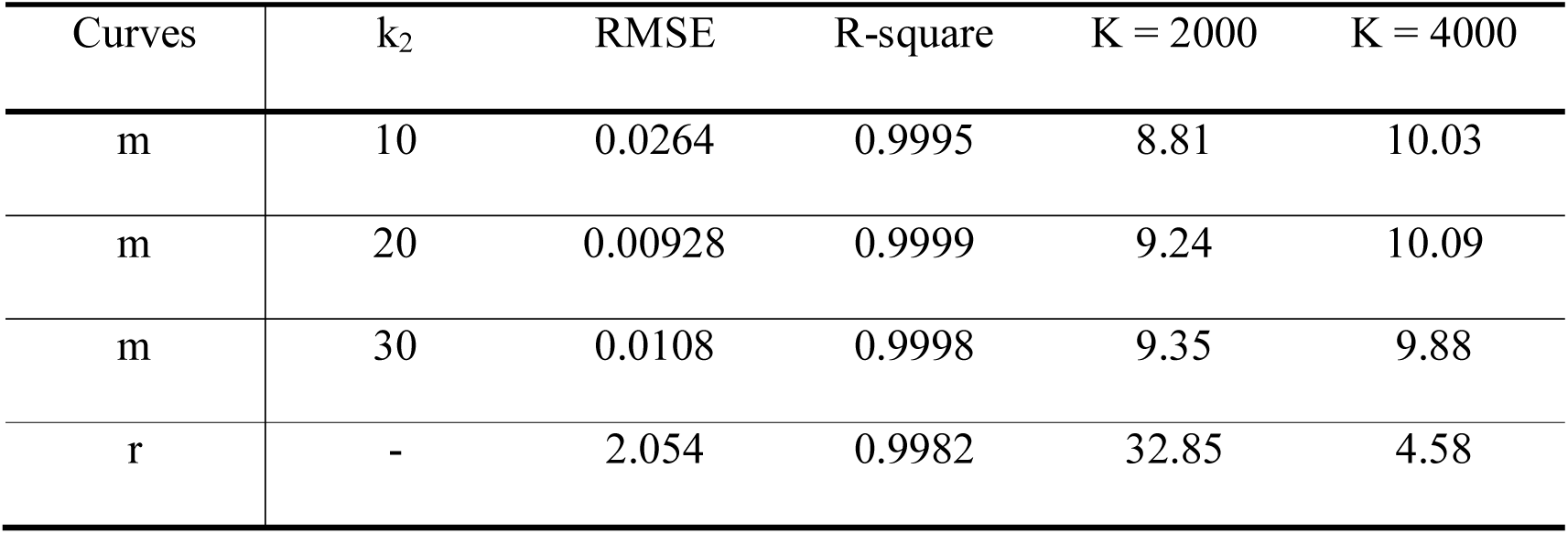
Performance of curve fitting and predictions.

| Curves | $k_2$ | RMSE | R-square | K = 2000 | K = 4000 |
| --- | --- | --- | --- | --- | --- |
| m | 10 | 0.0264 | 0.9995 | 8.81 | 10.03 |
| m | 20 | 0.00928 | 0.9999 | 9.24 | 10.09 |
| m | 30 | 0.0108 | 0.9998 | 9.35 | 9.88 |
| r | - | 2.054 | 0.9982 | 32.85 | 4.58 |

**Supplementary Figure 5:**
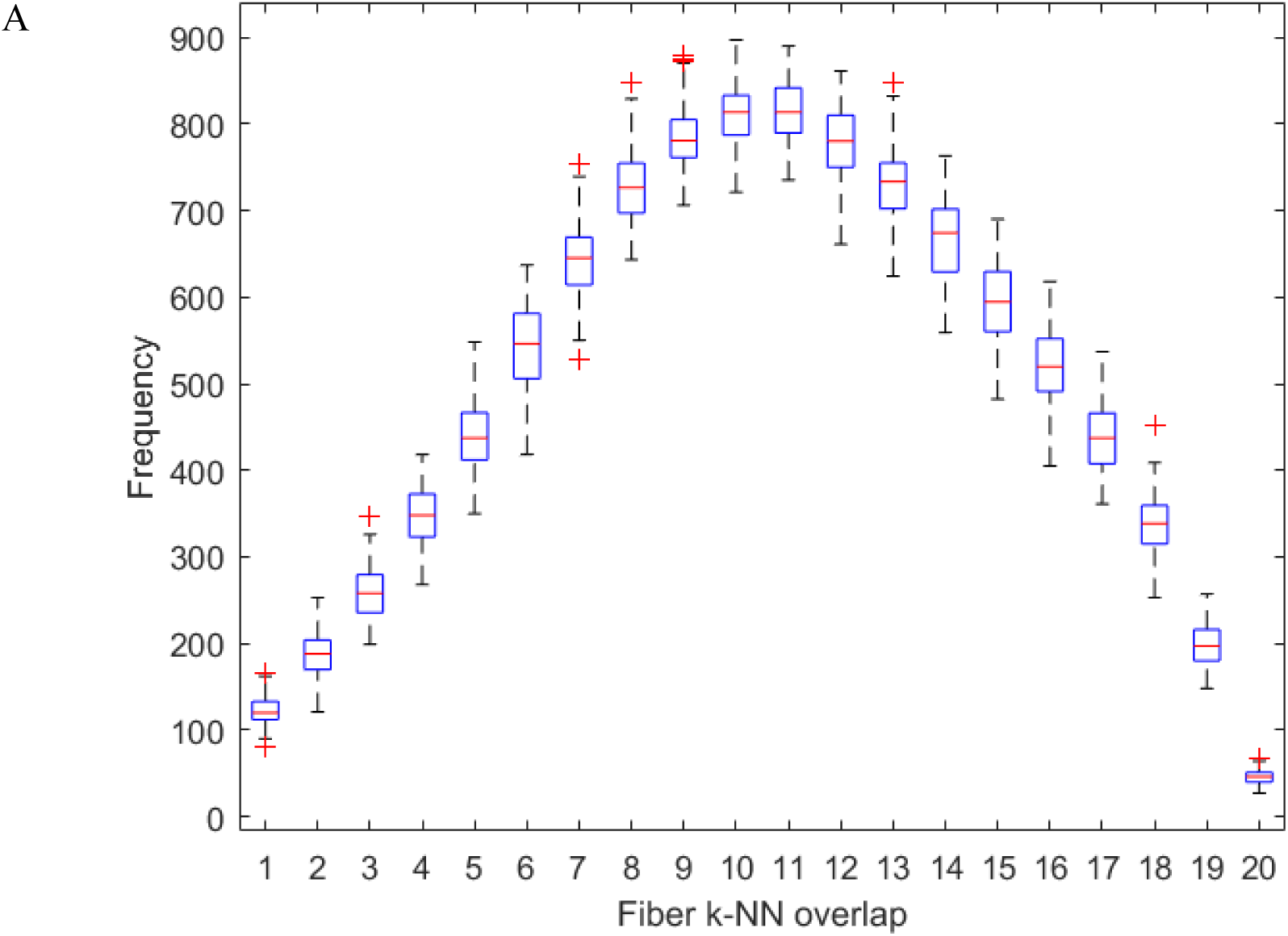
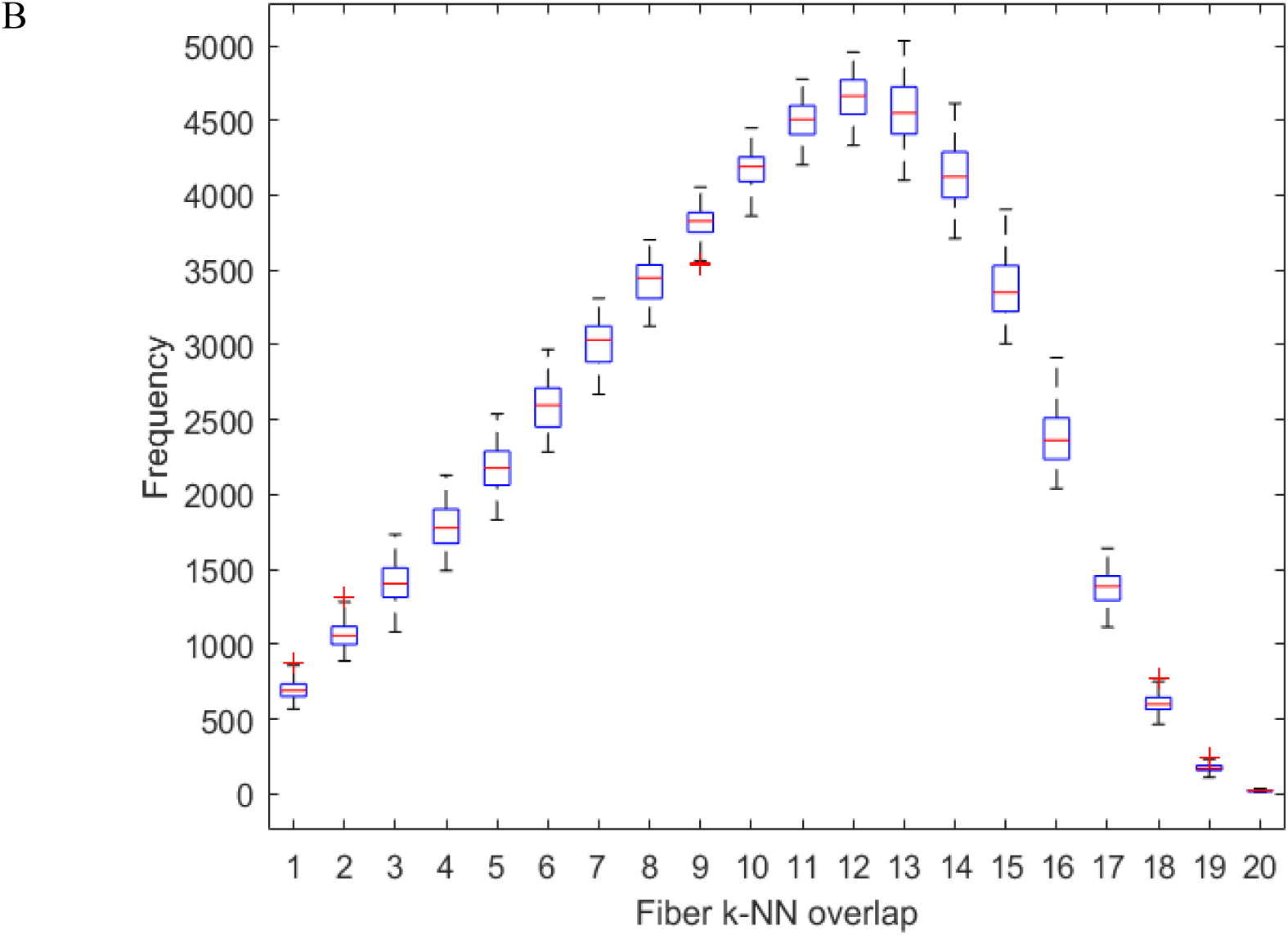
Performance of inter-subject fast streamline *k*-NN. A) and B) are the performance measures for tractograms with 10K and 50K streamlines^f^.

#### S3 Group-wise white matter topography analysis

##### MDS for white matter topography analysis

Our main idea for measuring streamline-wise white matter topography is based on classical Multidimensional Scaling (cMDS) ^21,22^.

###### Basic formulations of MDS

The cMDS embedding can be obtained by using the following steps:

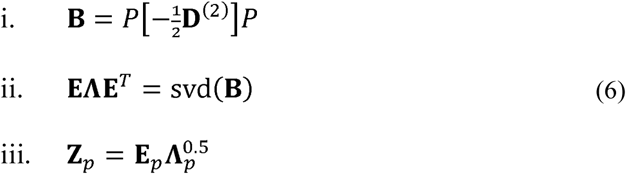

where *P* is known as the centering matrix defined as 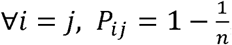, and ∀*i* ≠ *j*, 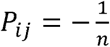, where *n* is the number of samples, **D**^(2)^ is the input squared pairwise distance matrix, **Z**_*p*_ and **E**_*p*_ are the first *p* columns of **Z** and **E**, and **Λ**_*p*_ is the top-left *p* × *p* block matrix of **Λ** where *p* is the embedding dimension. In this work, we adopt all the eigenvectors associated with positive eigenvalues for embedding.

###### The geometry of streamline streamlines in the embedding space

Based on the MDS formulation, we can infer that more contrast on the values of embedding vectors can be seen for geometrically more distant streamlines. In addition, the significance of each dimension of the embedding vectors in terms of its contribution in forming the target distance is naturally characterized by the corresponding eigenvalues. This can be seen from the maximum principle of Rayleigh quotient:

###### Theorem 2.

*Let A be Hermitian and suppose its eigenvalues are λ*_1_ ≤ *λ*_2_ … ≤, *λ*_*n*_, *and their associated eigenvectors are* ***e***_1_,***e***_2_,…,***e***_*n*_:

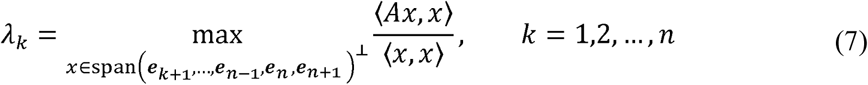

*and the maximum is attained for x =* ***e***_*k*_, *where* ***e***_*n*+1_ = **0.**

The proof is straightforward by recursively applying the maximal principle of Rayleigh quotient^62^ to the residual subspace of *A*. In words and in a loosen sense, the eigenvectors of *A* maximally correlate with *A*, and the correlation is quantified by the eigenvalues. In our case, *A* = **B** is the Gram matrix.

An alternative and more intuitive view of the decomposition is the following:

###### Proposition 3.

*Let* **B** *and* **z**_*i*_, *i* = 1,2,…*be the Gram matrix and the MDS embeddings defined in Eq. (6)*:

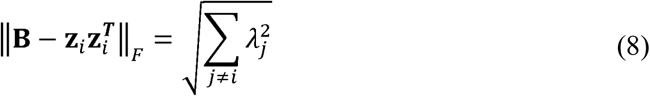

*where* **Λ** = diag(*λ*_1_,*λ*_2_,…) *are the eigenvalues associated with the eigenvectors* **E** = [**e**_1_, **e**_2_,…].

The proof is deferred to the appendix.

In conclusion, the embedding space coincides with the metric-distance geometry of the original streamlines, and each dimension represents a subspace of the geometry. Furthermore, they resemble the classic yet vague notion of topographic gradients in the neuroscience literature. Accordingly, we call them the *topographic vectors*.

##### Group-wise white matter topography analysis

To extend the individual white matter topography analysis to group-wise analysis, we will need to compare the embedding vectors for a group of subjects. Our idea is based on the conventional group-wise analysis of volumetric data in which the brain voxels are registered and then analysed across the group. To achieve this, we will need to address two technical challenges. First of all, the topographic vectors computed on parallel for each individual may be misaligned, largely due to brain symmetry and non-uniqueness of the MDS, i.e. 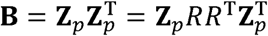 where *R*^*p*×*p*^ is an arbitrary orthogonal matrix. Secondly, a topographic vector of one subject does not correspond to any of the topographic vectors of another subject and the order of the topographic vectors is the same as the order of the streamline streamlines in the whole brain tractogram which is random for any subject.

The first problem renders the topographic vectors of different subjects incomparable, and the second problem prohibits us from comparing the comparable topographic vectors. Accordingly, we propose to align and match the topographic vectors across group of subjects. The alignment will recover the common *pose* of the topographic vectors by finding the aligning orthogonal matrix *R* for each subject. Afterwards, the topographic vector matching is used to further reduce the individual differences and identify the correspondences in the topographic vectors. A by-product of this work is that it achieves non-rigid whole-brain tractogram matching, which is by itself a well-known technical challenge to the neuroimage computing community.

###### Topographic vector alignment

Without loss of generality, here we consider the case for two subjects. Let the two streamline bundles be denoted as *T*_1_ and *T*_2_. Their topographic vectors are denoted as 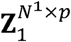 and 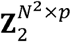. Similar to the volumetric studies where the alignment can be done using the similarity measure of voxels, we may attain the alignment using the similarity between the streamline streamlines across different subjects. However, as we have discussed previously, computing the pairwise streamline *k*-NN is time-consuming. Accordingly, we adopt the fast streamline *k*-NN instead. Suppose we consider *T*_1_ as a fixed reference, we can find the subset of **Z**_2_ corresponding to **Z**_1_ based on approximate streamline *k*-NNs computed by our the fast streamline *k*-NN algorithm, and we denote it as **Z**_21_. Then, our formulation for cross-subject topographic vector alignment reads:

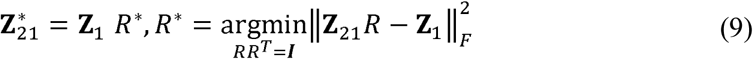

To solve this problem we first rewrite the objective function as

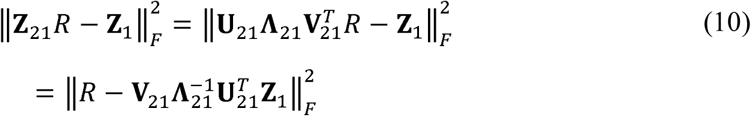

which is in the typical form of orthogonality-constrained least-square problem^63-65^, or it could be viewed as a variant of the Orthogonal Procrustes analysis^66^ and it admits a closed-form solution:

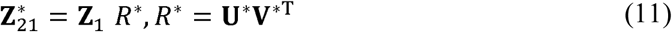

where

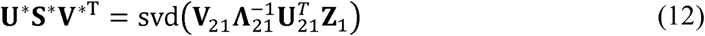

Note that this alignment does not change the within-tractogram pairwise distances, since the optimal transformation is orthogonal. The rationale of using an approximate *k*-NN rather than the exact *k*-NN is that the formulation is based on least-square cost which is robust to random errors.

The transformation may also be solved by using *normal equation* without imposing the orthogonality, and the pairwise distances in the aligned topographic vectors are not necessarily preserved. This choice is not theoretically sound but it may be more effective for initializing topographic vector matching. In terms of alignment, Eq. (11) is more reliable than normal equation.

###### Topographic vector matching

Upon finishing the global alignment of the topographic vectors, we proceed to establish the correspondences of the topographic vectors while refining the alignment to further remove the remaining minor misalignment due to the random errors in the global alignment, tractography and dMRI measurement. To this end, we cast the problem of whole-brain topographic vectors matching as a *point set registration* problem. Specifically, we propose to iteratively register the high dimensional topographic vectors across the different topographic vector point sets based on the iterative-closest-point criterion ^67,68^. This registration procedure automatically produces the point correspondences and alignment. Particularly, we adopt linear transformation in the point-set registration. Since all tractograms have been warped to a common space beforehand, the majority of the variability has been removed, and we consider the linear transformation to be sufficient to compensate for the remaining misalignment and random errors.

The iterative-closest point registration is known to be sensitive to initialization and can be easily stuck to local optimal solutions. Hence, the global alignment described before is used as the initialization of this step. We summarize our method for the group-wise topographic vector analysis in Algorithm 1.

###### Algorithm 2: Group-wise topographic vector analysis

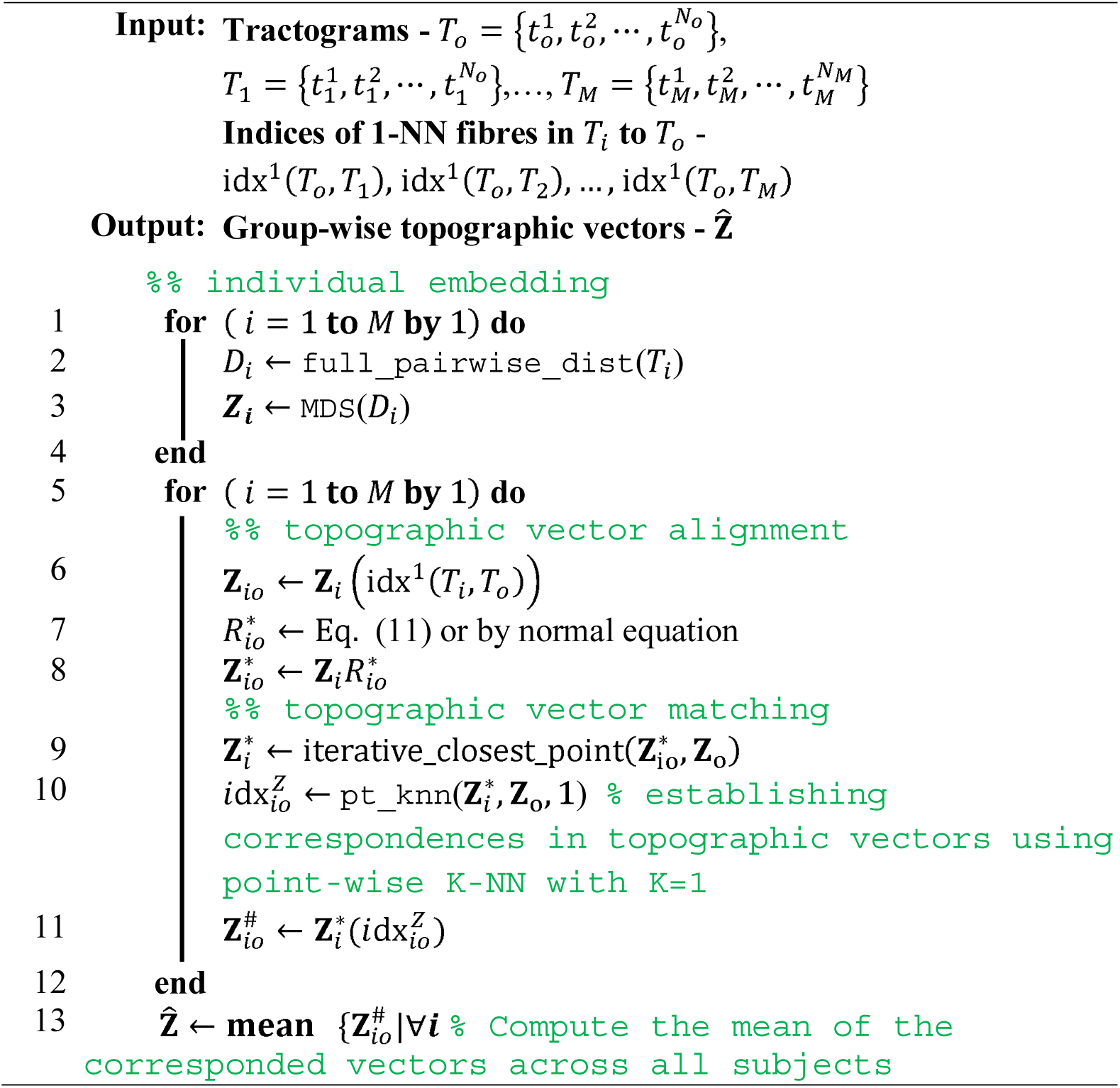

In terms of computational complexity, the point cloud registration is much simpler than direct streamline-wise matching or registration. Hence, this method can also be used for tractogram matching and registration.

###### Reliability of group-wise topographic vector analysis

Here, we present our essential performance assessment of our method for the topographic vector alignment and matching and the group-wise topographic vector analysis. In this experiment, we generated 100 pairs of whole-brain tractograms: one in the subject native T1 space, the other is warped to 100307 T1 native space using TDI volumetric registration with ANTs^46^. Each tractogram contains 10K streamline curves. We then evaluate our method by matching the ones in subject native space to the ones in the warped space.

We compare our method against the original topographic vectors without registration, and we also compare our method with the topographic vector registration without initialization using fast streamline *k*-NN algorithm (FSkNN). Some visual results of the linear iterative-closest-point (ICP) based topographic vector registration are shown in **Supplementary Figure 6**. Our results with initialization by linear transformation and orthogonal transformation match visually very well to the reference, while the matching without initialization by FSkNN algorithm can easily be stuck to local optimal solutions. The results with initialization by linear transformation are slightly better than the results with initialization by orthogonal transformation, although the orthogonal transformation preserves the pairwise distances and it is theoretically more sound.

Since the correspondences between the streamlines in the pairs of tractograms are known in this experiment, we are able to quantitatively evaluate the performance of the streamline matching. The histograms of the matching accuracy of different methods and settings are shown in **Supplementary Figure 7**. The matching accuracy is defined as the 1-average Hamming distance, or 1 - average matching error, between the ground-truth matching correspondences and the results. We further present the statistics of the matching in **Supplementary Table 2**. From both the histograms and statistics we observe that our method dramatically improves the streamline matching performance over the matching without initialization by FSkNN algorithm.

The performance of our FSkNN distance computation for group-wise white matter topography analysis is quantitatively assessed by comparing the aligned topographic vectors computed via our FSkNN algorithm with the ones computed with exhaustive streamline distance computation. We used three measures to quantify the performance: a) element-wise Pearson linear correlation of all topographic vectors denoted as `corr` b) mean RMSE (M-RMSE) of the topographic vectors overall dimensions. We present the statistics in **Supplementary Table 3**, according to which we can conclude that our method produces topographic vectors almost identical to the ones given by exhaustive streamline distance computation in terms of linear correlation.

Note that the range of each dimension of the topographic vectors is in tens to hundreds. The absolute difference between the results from the two distance computation methods is also negligibly small. Furthermore, alignments with both of the two distances achieve high fidelity, in terms of M-RMSE, and reproducibility, in terms of the standard deviation of RMSE, when comparing the alignment results with the reference data.

**Supplementary Figure 6:**
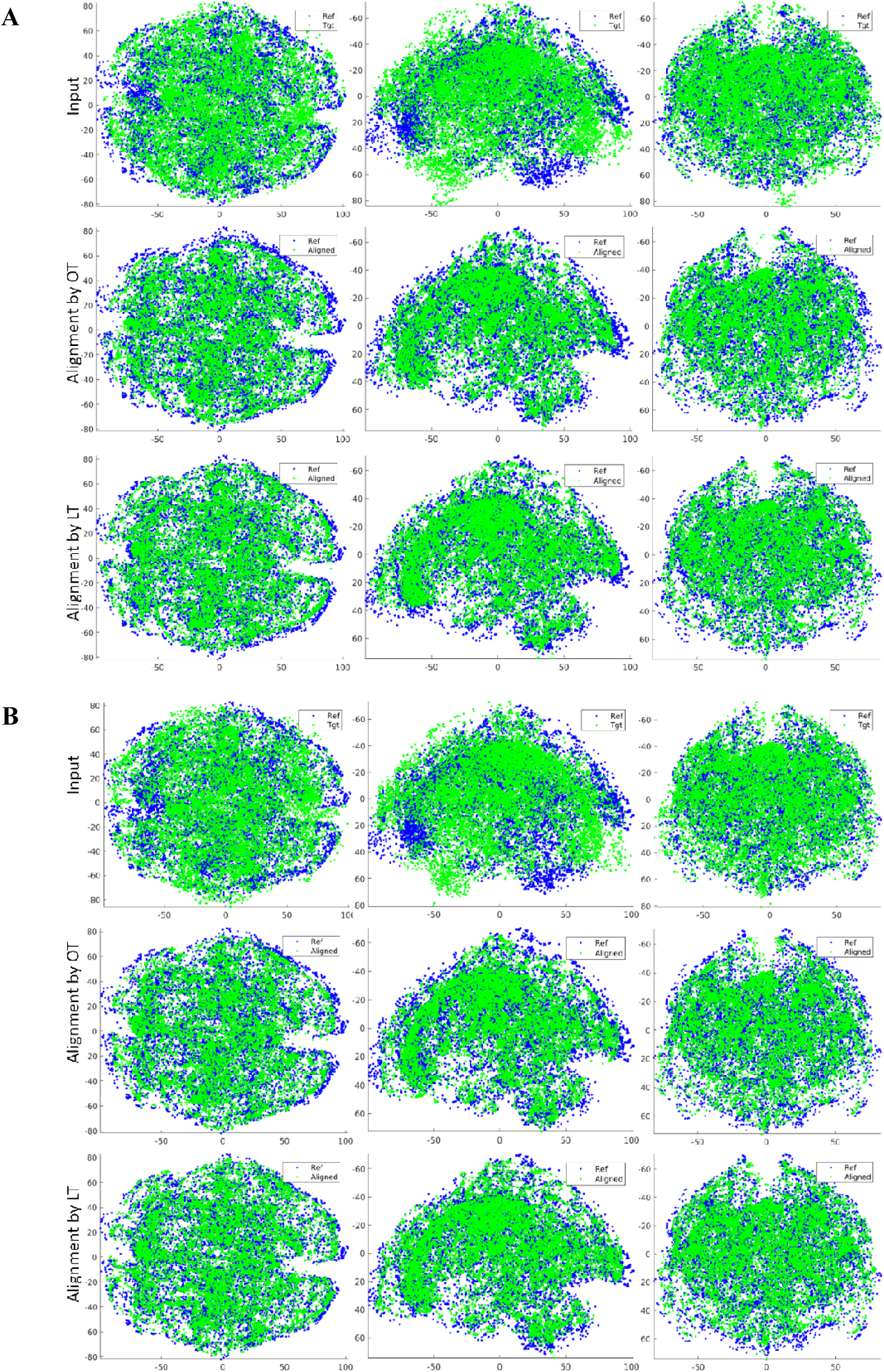
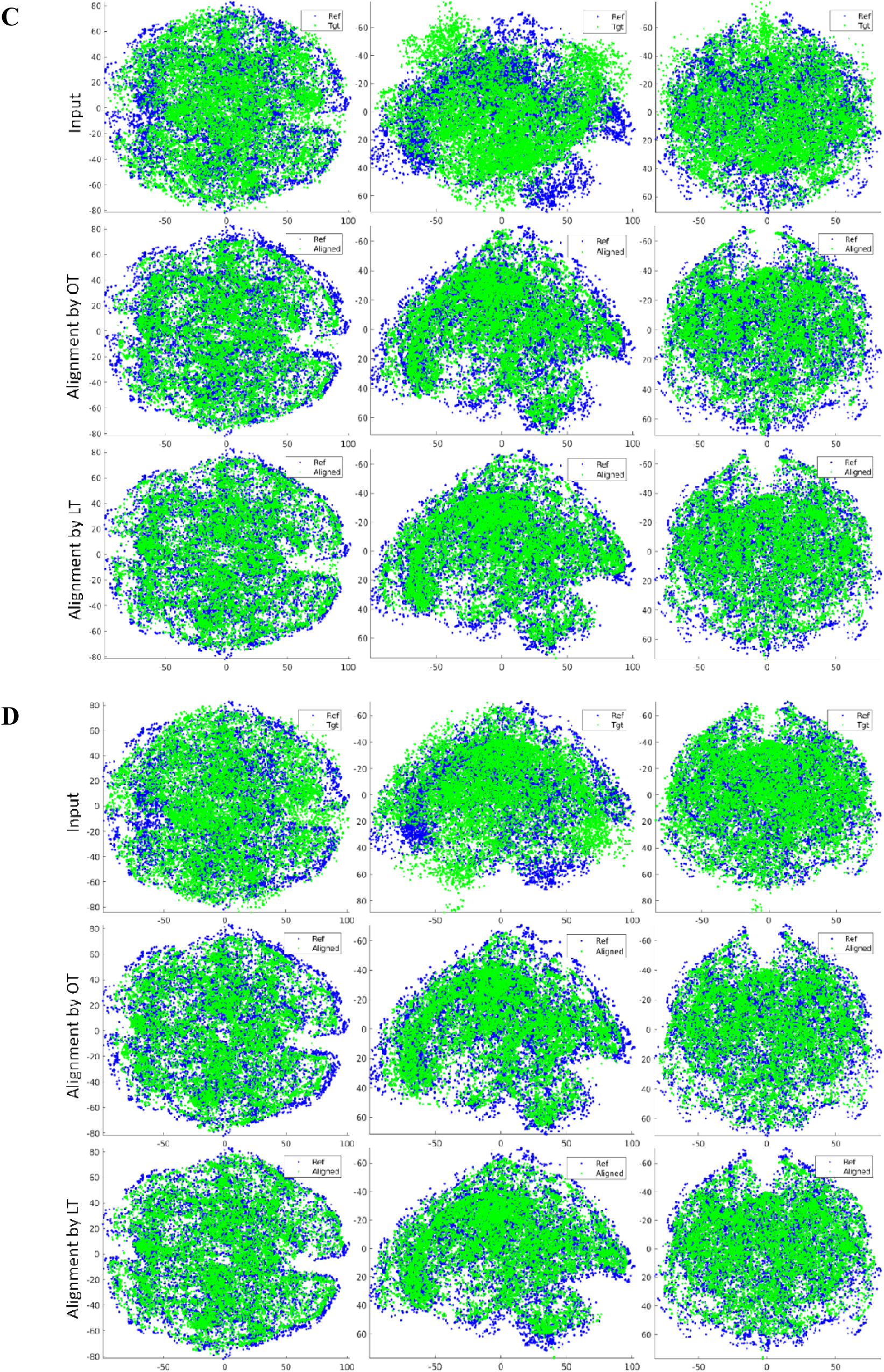
Effect of topographic vector alignment for HCP subjects. **Ref**: the topographic vectors of reference subject. **Tgt**: the topographic vectors of the target subject. **Align**: the aligned topographic vectors of the target subject. **OT** is the result of alignment with initialization by orthogonal transformation. **LT** is the result of alignment with initialization by linear transformation.

**Supplementary Table 2:** Matching performance.

|  | GT vs Input | GT vs Aligned<br>w/o FSkNN | GT vs Aligned<br>with FSkNN + OT | GT vs Aligned<br>with FSkNN + LT |
| --- | --- | --- | --- | --- |
| **Accuracy** | 0.0146 | 0.1950 | 0.8973 | 0.9051 |

**Supplementary Figure 7:**
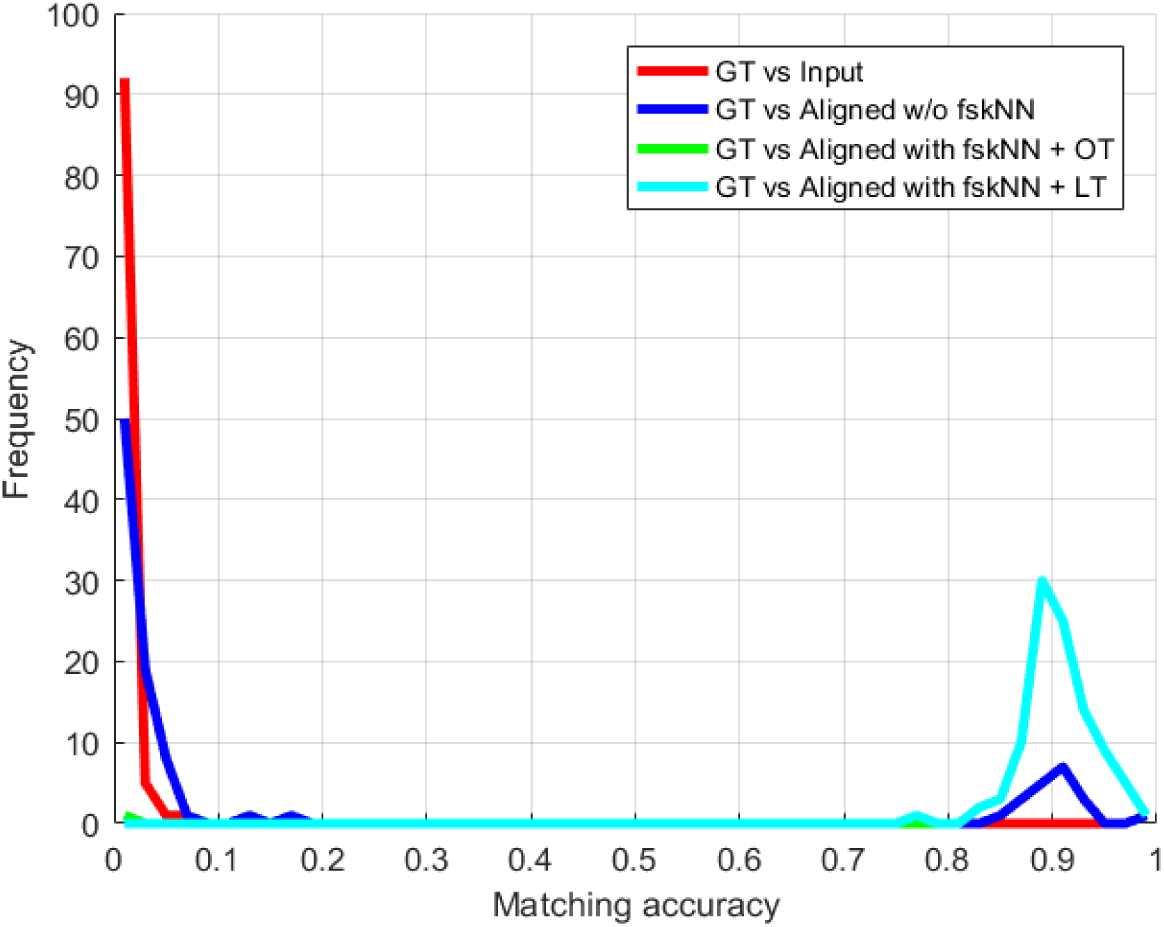
Performance of non-rigid whole-brain tractogram matching. a) Visualization of tractogram matching in embedding space. b) Histograms of average matching accuracy.

**Supplementary Table 3:**
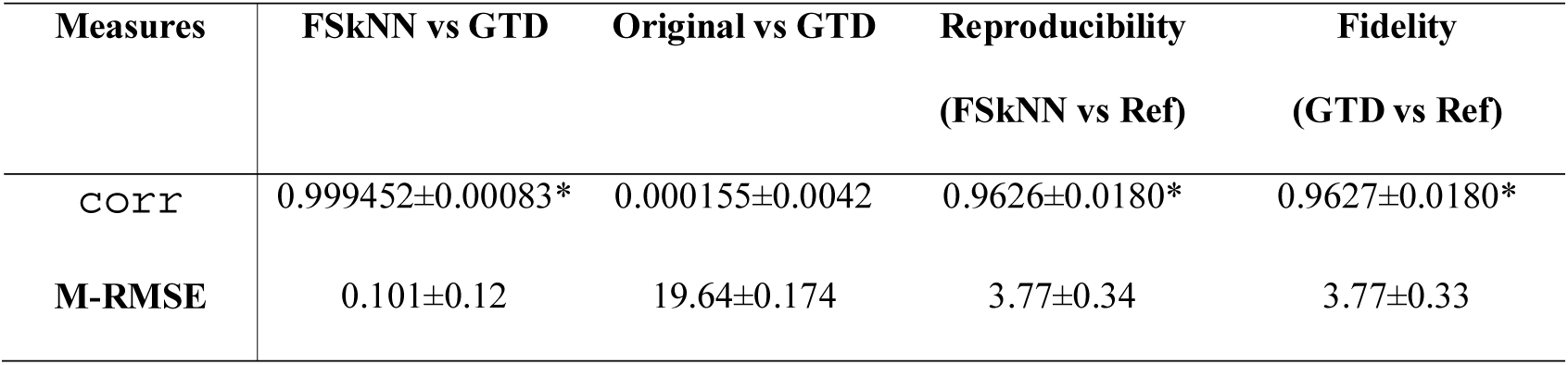
Alignment and matching performance (* indicates statistical significance at α=0.001)

###### Additional results of group-wise topographic vector analysis

In **Supplementary Figure 8**, we show the two-dimensional histograms of the top-10 group-wise topographic vectors against the streamline coordinates. The strongly linear correlation is only observable in the top-3 dimensions of the topographic vectors. In the remaining dimensions, certain correlation can be observed, and the histograms may form regular patterns. Non-trivial correlation values can be found in some of the remaining dimension, e.g. 4^th^, 5^th^ and 9^th^.

**Supplementary Figure 9:**
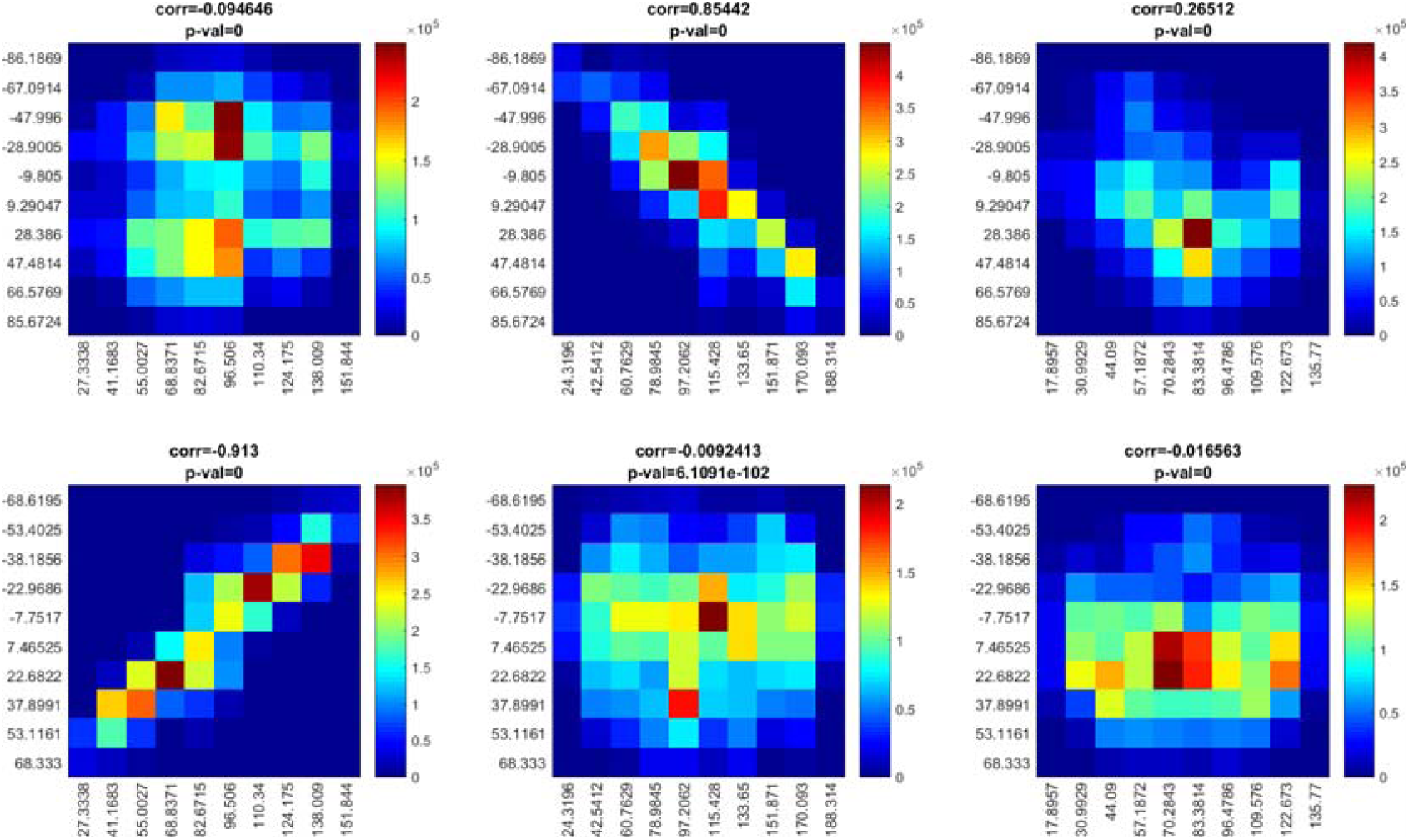
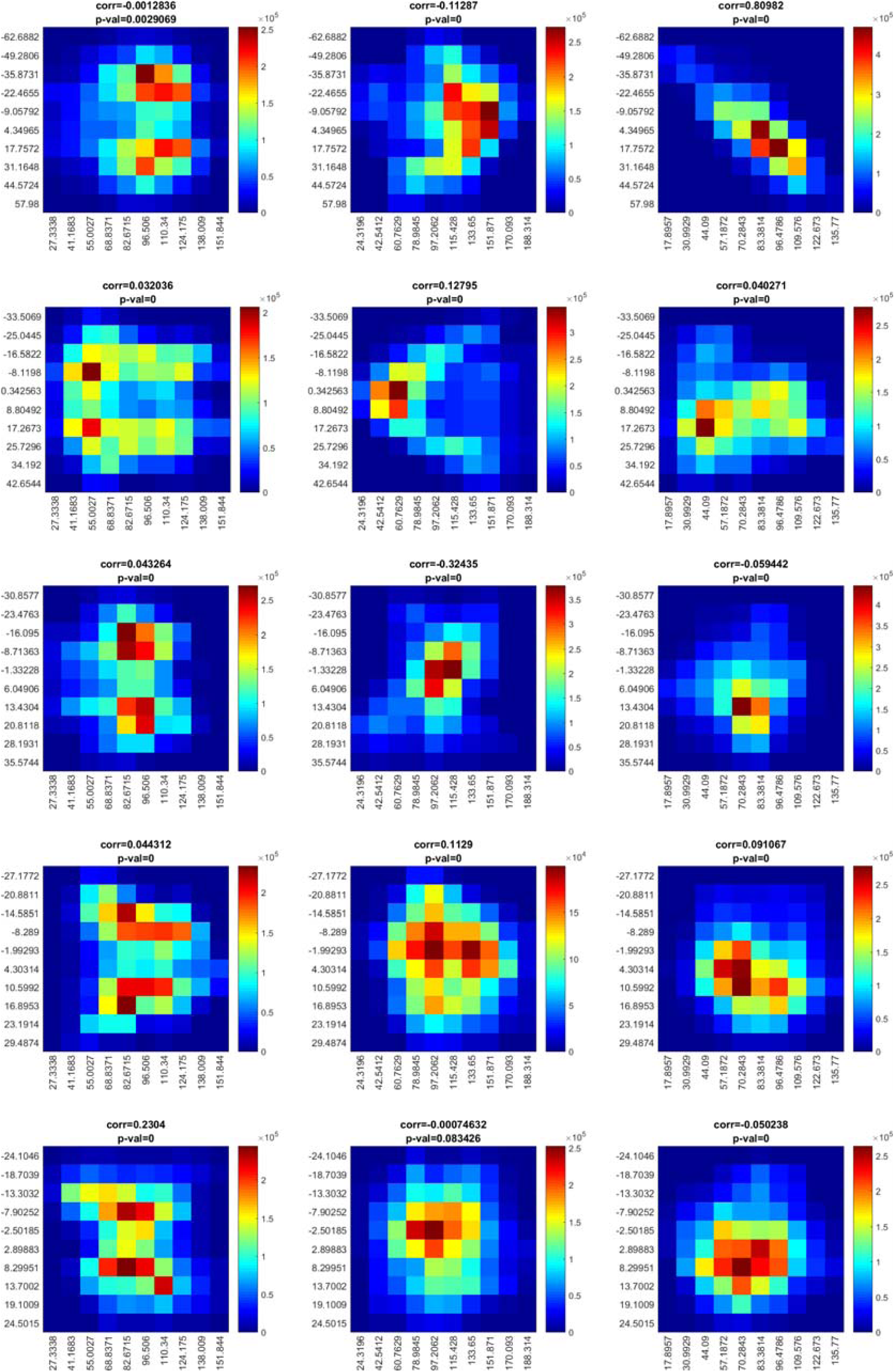
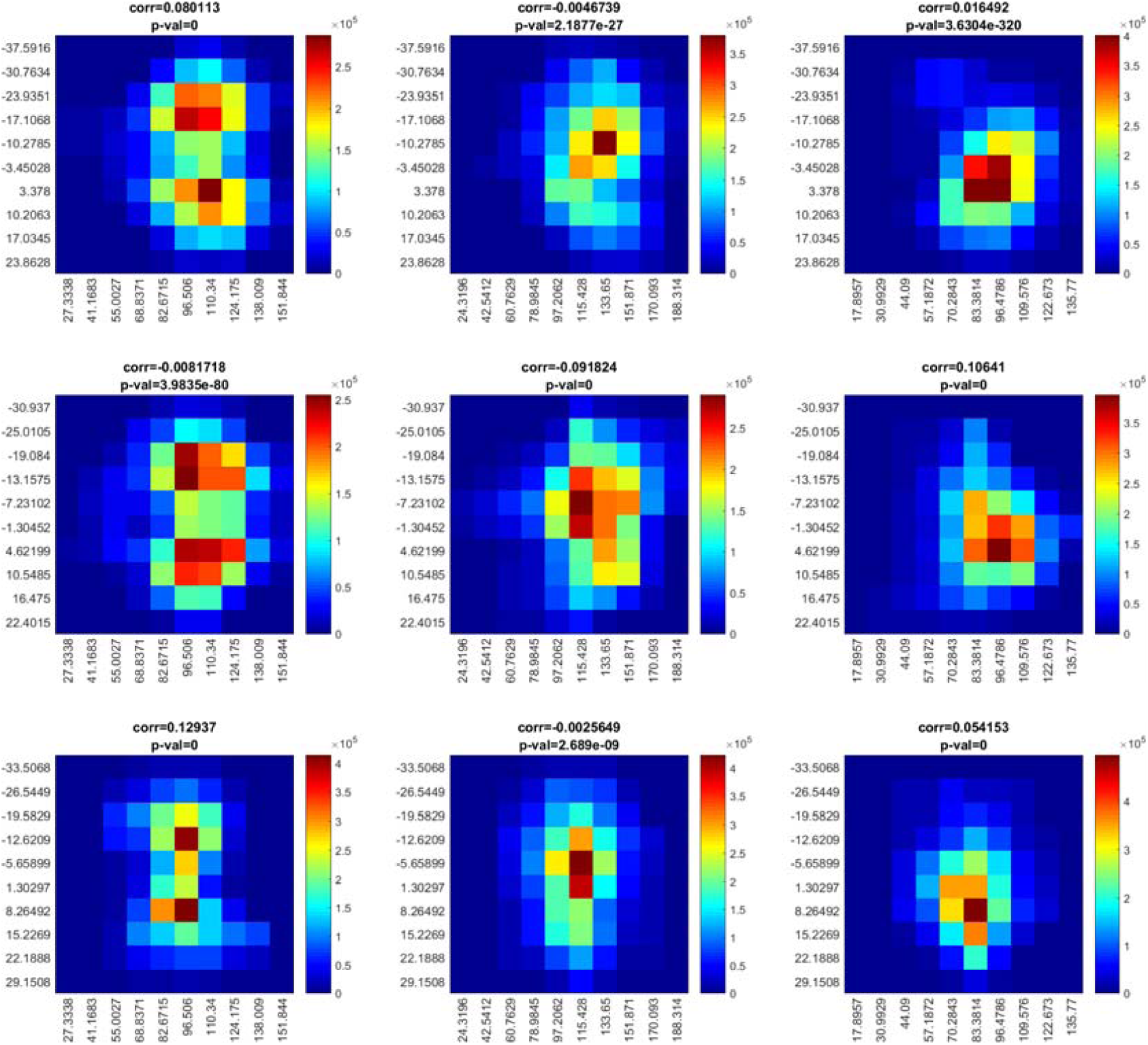
From the top row to the bottom row are the two-dimensional histograms of the 1^st^ to 10^th^ dimensions of the topographic vectors against the streamline coordinates. The correlations and p-values are displayed on top of each histogram.

## Appendix

### Proof of Theorem 1

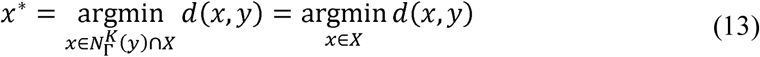

Since 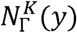 is the set of point-wise K-nearest points of *y* in Γand Γcontains point set *X*, and since 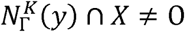, we can define two subsets of *X* as *X*_*y*_ and 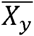 satisfying 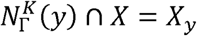 and 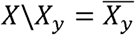. Based on the definition of the point-wise K-nearest neighbour 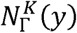, we know that

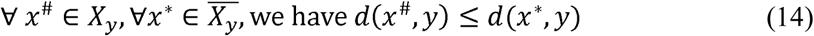

Therefore,

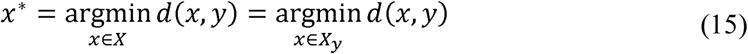

which completes the proof. ▪

**Proof of** Error! Reference source not found.

The L.H.S. of Eq. (X) can be rewritten as

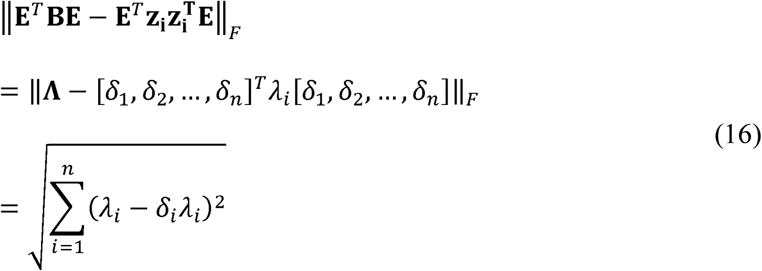

which completes the proof. ▪

## Footnotes

a http://trackvis.org/

b Unless otherwise stated, the central mark indicates the median, and the bottom and top edges of the box indicate the 25th and 75th percentiles, respectively. The whiskers extend to the most extreme data points not considered outliers, and the outliers are plotted individually using the ‘+’ symbol.

c Code downloaded from https://www.mathworks.com/matlabcentral/fileexchange/58312-kernel-density-estimator-for-high-dimensions

d In the boxplots of (C) and (D), all the parameters are defined as the same as in other boxplots, except that the whiskers spread over the entire span of the data distribution, and the outliers are no longer visualised for simplicity of the presentation.

e A multiset is a modification of the concept of a set that, unlike a set, allows for multiple instances for each of its elements.

f The central mark indicates the median, and the bottom and top edges of the box indicate the 25th and 75th percentiles, respectively. The whiskers extend to the most extreme data points not considered outliers, and the outliers are plotted individually using the ‘+’ symbol.

